# EcoCore; An ecologically diverse panel of *Arabidopsis thaliana* accessions for studying plant-environment interactions

**DOI:** 10.64898/2026.04.17.719158

**Authors:** Abraham L. van Eijnatten, Jelle J. Keijzer, Jana Trenner, Carolin Delker, Marcel Quint, Martijn van Zanten, L. Basten Snoek

**Affiliations:** Theoretical Biology and Bioinformatics, Institute of Biodynamics and Biocomplexity, Utrecht University, 3584 CH Utrecht, The Netherlands; Plant Stress Resilience, Institute of Environmental Biology, Utrecht University, 3584 CH Utrecht, The Netherlands; CropXR Institute, Utrecht, the Netherlands; Institute of Agricultural and Nutritional Sciences, Martin Luther University Halle-Wittenberg, 06120 Halle (Saale), Germany; German Centre for Integrative Biodiversity Research (iDiv), Halle-Jena-Leipzig, Germany

## Abstract

*Arabidopsis thaliana* naturally occurs across a wide geographic range and displays extensive natural variation in several traits including adaptive responses to the abiotic environment (e.g. temperature, drought, salt). Quantitative techniques like Genome Wide Association Studies (GWAS) enable mapping the genetic basis of such environmental responses and benefits from extensive genetic variation, but the size of the chosen diversity panel is often limited by phenotyping capacity. Most studies therefore use subpanels, often based on maximization of genetic diversity. However, this type of selection may overrepresent cosmopolitan alleles and underrepresent rare environment-specific alleles.

Here, we demonstrate that the genetic variation in a GWAS subpanel of *Arabidopsis thaliana* accessions depends almost entirely on the number of accessions in the panel and very little on the composition of the panel. We present the EcoCore panel designed by grouping accessions of the 1001 genomes (1001G; 1135 accessions) collection, based on their native collection environment and selecting an equal number of accessions from each environment. We assessed hypocotyl lengths of plants grown at control and ambient high temperatures (20°C and 28°C) for 913 accessions of the 1001G and mapped these traits with the full 1001G panel versus the EcoCore panel. The EcoCore panel revealed novel genetic associations with hypocotyl length which is attributed to enrichment of alleles from rare environments. We present the EcoCore panel as a manageable resource for studying phenotypic plasticity and the genetic basis of plant–environment interactions.

## Introduction

The plant species *Arabidopsis thaliana* has a very broad geographic distribution and is native to Eurasia and parts of Africa, spanning diverse ecological and environmental conditions (Koornneef et al., 2004; Yim et al., 2024). The geo-biographical distribution is limited by cold winter temperatures at northern latitudes, and by summer heat and elevation at southern latitudes (Hoffmann, 2002; Yim et al., 2024). Drought also puts a limit on Arabidopsis performance (Clauw et al., 2015), likely constraining the species’ distribution. As a pioneer species, Arabidopsis evolved many phenological adaptations and exhibits strong phenotypic plasticity to cope with local biotic and abiotic environmental conditions, for example temperature (Ibañez et al., 2017; Jiang et al., 2024, 2025; Morales et al., 2022; Quint et al., 2016; Scheepens et al., 2018; Vasseur et al., 2018). This resulted in broad natural variation in traits mediating environmental responsiveness (Koornneef et al., 2004). For instance, Arabidopsis plants from colder climates are often relatively cold tolerant, have an insulating compact architecture (Hopkins et al., 2008), and require a prolonged period of cold (vernalization) to induce floral competence in spring (Johanson et al., 2000; Lee et al., 1994; Sung & Amasino, 2004). In contrast, accessions from warmer climates show signs of adaptation to high temperatures (Ågren & Schemske, 2012; Li et al., 2010; Tonsor et al., 2008; Wolfe & Tonsor, 2014). Notably, warmth-exposed Arabidopsis plants exhibit an open rosette architecture (Ibañez et al., 2017), and this phenotype scales with latitude of occurrence to optimize photosynthetic output caused by differences in light intensities (Hopkins et al., 2008). Additionally, Arabidopsis plants occurring in warm environments may adopt a spring-annual life cycle wherein the seeds germinate in spring and flowering and seed set takes place in late spring/early summer (Donohue et al., 2008; Montesinos‐Navarro et al., 2011; Scheepens et al., 2018).

Studying environmental plasticity and its genetic basis using *Arabidopsis thaliana* as a model can help improve crop species’ performance to ensure food security in the context of climate change (Brady et al., 2025; Roeder et al., 2025). The genetic basis of environmental plasticity can be elucidated by quantitative genetic techniques such as Genome-Wide Association Studies (GWAS). GWAS relies on associating standing natural genetic variation, such as single nucleotide polymorphisms (SNPs), to trait values in a population, and can be used to elucidate genomic features underlying local adaptation to climate (Exposito-Alonso et al., 2019; Ferrero-Serrano & Assmann, 2019; Li et al., 2010). For effective GWAS analysis large diversity panels of accessions need to be genotyped for genetic variation and phenotyped for the trait(s) under study. Several fully-sequenced diversity panels are publicly available, such as the ‘1001 Genome (1001G) panel’, which consists of 1135 high quality genome-sequenced near homozygous natural Arabidopsis accessions derived from across the species’ distribution range (The 1001 Genomes Consortium et al., 2016; The 1001 Genomes Plus Consortium et al., 2024). Such diversity panels provide a detailed map of standing genetic variation that is readily available for mapping, making phenotyping the major bottleneck for GWAS.

Despite the increasing availability of high-throughput phenotyping power (Yang et al., 2020; Zavafer et al., 2023), logistical, financial, and time constraints can still pose limits on the number of accessions that can be realistically phenotyped for quantitative genetic approaches. To overcome this bottleneck, researchers often use sub-sets of diversity panels (subpanels) (Seren et al., 2017), whereby selection is usually based on maximizing the genetic diversity in the subpanel (Li et al., 2010; McKhann et al., 2004). Although meaningful, this approach potentially enriches for accessions of ‘common’ environments, whereby SNPs causal for specific adaptations of relatively unique environments are lost due to their low frequency in the subpanel (Floriani & Lipka, 2025). Thus, we argue that studies exploring the role of genetics in environment-imposed traits, could benefit from maximizing variation in environmental parameters, instead of genetic variation.

In this work we present a core-set of 289 natural Arabidopsis accessions, derived from the 1135 genotypes of the full 1001G panel, that exhibits optimized variance in environmental and ecological parameters defining the collection site balanced with ample genetic variation. We dub this subpanel the ‘EcoCore panel’. To assess the effect of selecting accessions from maximally different environments on genetic mapping studies, we assessed the phenotypic diversity in hypocotyl lengths for almost the full 1001G panel and compared GWAS results between the full 1001G panel and the EcoCore panel. Our data indicate that the EcoCore panel can enrich rare and potentially relevant alleles from specific environments that are underrepresented in the 1001G panel, thus revealing novel associations that remain hidden when assessing the full panel. We put the EcoCore panel forward as an effective and manageable-sized research collection aimed at assessing the genetic basis of environment-imposed traits.

## Results

### Genetic diversity scales with panel size, not composition

The experimental setup for natural variation studies is often resource-limited by the capacity to phenotype large numbers of accessions. In our attempt to construct a feasibly-sized subpanel of the 1001G optimized for studying environment-imposed traits, we therefore started with questioning what panel size is deemed feasible. To address this, we used the AraPheno database (Seren et al., 2017; Togninalli et al., 2019) and a curated extension of this database (Ruffley et al., 2023) to probe typically used panel sizes. The median subpanel size used was 135 accessions for the AraPheno database (**Supplemental Figure 1, Supplemental Dataset 1**; corresponding to ∼11% of the 1135 accessions of the 1001G panel) and 102 accessions per trait for the extended database (**Supplemental Figure 2**). Some studies in the AraPheno database used panels of which part of the accessions are not included in the 1001G panel, such as the HapMap panel (Li et al., 2010). Measurements corresponding to these accessions were excluded from the AraPheno database, thus leading to an underestimation of the actual panel sizes. Correcting this aberrancy by extracting the used panel size from the source literature (**Figure 1A, Supplemental Dataset 2**) indicated that the median panel size per study was 310 accessions (corresponding to ∼27% of the 1001G panel).

**Figure 1:**
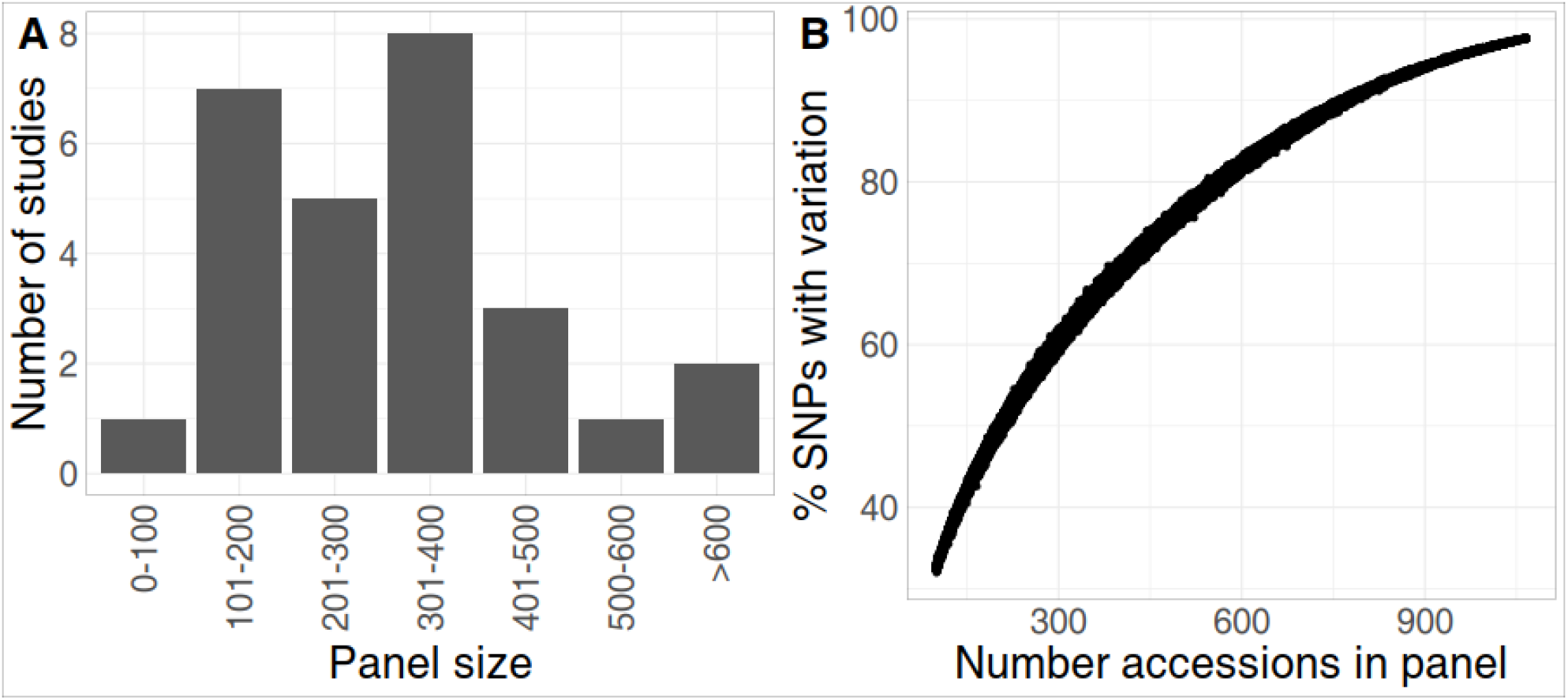
Genetic variation scales with panel size. **A)** Number of studies in the AraPheno database in panel size categories. The panel sizes were obtained from the original publications. **B)** Relation between SNP variation and number of accessions used. Indicated are the fraction of SNPs of the full 1001G set with considerable variation (at least 13 accessions with the reference and 13 with the alternate allele) in random sub-panels of the full 1001G panel (100 random subpanels generated for each panel size between 100 to 1065 accessions).

Next, we assessed for each accession of the 1001G the number of studies in which it was included in the phenotyped panel. This revealed a median of 4 studies per accession, with only 62 accessions assessed in more than 15 studies (**Supplemental Figure 3A, Supplemental Dataset 3**). The accession that was included in the most studies was Tottarp-2 (22 out of 25 studies) followed by Col-0, An-1, Bor-4, Mz-0, and Aa-0 (all 21 studies). The number of studies in which accessions were included depended on the admixture group of the accessions (obtained from The 1001 Genomes Consortium., 2016), with accessions from Germany and south-Sweden admixture groups appearing in the most studies (**Supplemental Figure 3B**). In summary, working with a sub-panel from the 1001G diversity panel of ∼300 accessions is common practice, and the used subpanels are biased towards specific accessions/admixture groups. Furthermore, the use of (close to) the full 1001G panel is rarely reported in literature thus far.

The subpanel composition may determine if certain genetic variants have sufficient variation for use in GWAS and may lead to bias towards a certain phenotypic range. To understand how panel composition is usually determined, we examined the justifications of panel selection of the studies incorporated into the AraPheno database. Some studies report the use of available subpanels, such as the HapMap panel (Li et al., 2010) (n = 7) or the Busch group collection (Ristova et al., 2016) (n = 2). The HapMap panel was constructed as a subpanel of a larger panel with 5810 accessions by maximizing genetic diversity, although the exact panel selection procedure was not revealed (Li et al., 2010). The Busch group collection covered the genetic diversity and geographic space of the 1001G panel, but did not explicitly maximize it (Ristova et al., 2016). Other studies in the AraPheno database made their own subpanels of the 1001G panel, mentioning that their subpanel is genetically or geographically diverse, but without mentioning concrete selection criteria (n = 5). Finally, some studies selected accessions from specific geographic locations to study local adaptation (n = 4), e.g. using only Swedish accessions.

Most studies thus used a subpanel selected for maximized genetic diversity. This approach assumes that the frequency of genetic variants/alleles in the subpanel needed for meaningful detection by GWAS can be substantially influenced by the composition of the subpanel. To investigate whether this assumption is correct we used a high-density genetic map (Arouisse et al., 2020) and generated 100 random subpanels of the 1001G panel of different sizes (100 – 1065 accessions, in total 96500 random panels) and plotted the number of genetic variants with considerable variation (Minor Allele Frequency (MAF) >= 13) against sub panel size (**Figure 1B**). This indicated that the number of alleles with considerable variation strongly scales with subpanel size, but is almost entirely independent of the panel composition. Hence, for quantitative genetic studies large(r) panels are obviously preferred because this maximizes the number of genetic variants available for GWAS. However, aiming to maximize the available genetic variants at a given subpanel size brings only small potential benefit. We also conclude that since the subpanel composition does not substantially matter for available genetic variants, it is valid to consider other selection criteria when designing a subpanel.

### Establishment of the EcoCore panel by maximizing environmental variation

Knowing the average size of subpanels used and that selection criteria other than maximized genetic diversity may benefit subpanel selection without losing genetic variation, we aimed to design a subpanel of the 1001G particularly suitable for studying environmental-responsive traits. Put in other words, our goal was to compose a balanced subpanel based on equal representation of accessions from diverse environments across the distribution range of *Arabidopsis thaliana* to ensure that sufficient genetic variation from each environment is present in the subpanel, even from rare environments.

To obtain a description of the environment at the local collection site of the accessions in the 1001G panel, we downloaded the AraClim database (Ferrero-Serrano & Assmann, 2019). This database contains 373 environmental parameters available for most of the 1001G (1066 accessions, **Supplemental Dataset 4**). Many of the 373 variables are descriptions of temperature and precipitation obtained by satellites, such as the frequently used WorldClim 2 parameters (Fick & Hijmans, 2017), while others are descriptors of e.g. soil properties, vegetation cover and greenhouse gas emissions. Principle Component Analysis (PCA) indicated that most of the variation between the collection sites can be attributed to temperature (PC1, explaining 34.1%) and precipitation (PC2, explaining 23.7%) parameters (**Supplemental Figure 4**). Because several of the 373 variables in AraClim are highly redundant, as shown by their correlation structure (**Supplemental Figure 5A**), we aimed to reduce the dimensions of the set by clustering the AraClim environmental parameters using k-means (K = 11, based on visual inspection of AraClim correlation network (**Supplemental Figure 5A**)) and used the clustering to select 11 non-redundant environmental parameters. These 11 non-redundant variables were descriptors of temperature, precipitation, humidity, and predicted GHG emission (**Supplemental Table 1**). Correlation network (**Supplemental Figure 5A**) and t-distributed stochastic neighbour embedding (t-SNE) analysis (**Supplemental Figure 5B**) showed that the 11 non-redundant variables were representative of the environmental parameters in the AraClim database. Furthermore, the 1001G accessions showed considerable clustering based on correlations between the 11 non-redundant environmental parameters (**Supplemental Figure 6**), indicating large variation in these parameters.

We next aimed to understand which 1001G accessions experience similar conditions in their local environment. To this aim, a hierarchical clustering was performed based on correlations between the 11 non-redundant environmental variables, to assign the 1001G panel accessions to 20 environmental groups (**Supplemental Figure 7, Supplemental Dataset 5**). Each environmental group shared a distinct local environment in terms of both the WorldClim 2 parameters (**Figure 2A**) and the 11 non-redundant variables (**Supplemental Figure 8**). Some environmental groups were geographically clustered, such as environmental group #17 which only occurs in central Germany, whereas other environmental groups covered a larger geographic range, such as environmental group #11, which occurred throughout central and eastern Europe, Russia, and in the north-west of the USA (**Supplemental Figure 9, Supplemental Figure 10**). The environmental groups also differ strongly in the number of included accessions (**Supplemental Figure 11**). For example, some environmental groups contained more than 100 accessions (environmental group #14; 142 accessions from Sweden and Finland, group #1; 114 accessions; predominantly from the center of the Iberian Peninsula, and group #6 (102 accessions predominantly from the south of Germany/ north of Italy). On the other end of the spectrum, some environmental groups contained less than 15 accessions (environmental group #12; 14 accessions, from the coast of Netherlands, north Germany, Japan, Sweden and Denmark), environmental group #16; 13 accessions, predominantly from Germany and Kyrgyzstan, environmental group #17; 12 accessions, all from Germany, and environmental group #18; 10 accessions, predominantly from Germany). We note that Germany stands out as it contains specific environments with little representative accessions in the 1001G, while accessions from environments in the center of the Iberian Peninsula and the south of Sweden are clearly overrepresented in the 1001G full panel.

**Figure 2:**
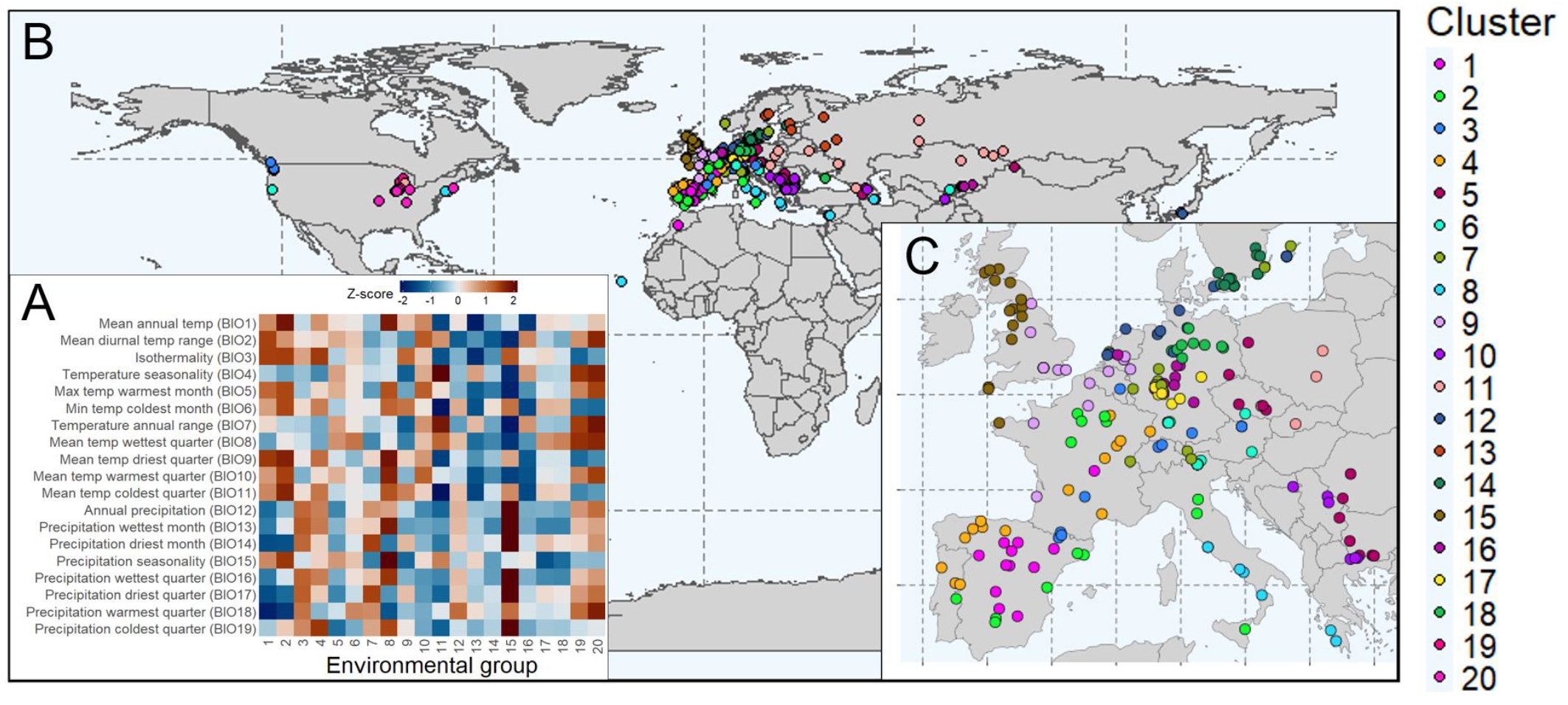
Global distribution of EcoCore panel accessions per environmental group. **A)** Heatmap showing for each environmental group the relative value expressed as z-score for WorldClim 2 variables obtained from the AraClim database. **B)** World map showing the geographic origin of accessions included in the EcoCore panel color-coded per environmental group. **C)** Zoom in on B), showing the geographic origins of the EcoCore panel accessions within Europe.

To construct a subpanel of ∼300 accessions (in line with the median panel size of 310 accessions for the studies included in the AraPheno database) covering maximally diverse environments, we subsequently selected 15 accessions from each of the 20 environmental groups based on maximized genetic dissimilarity within each environmental group. If there were 15 or less representatives, all accessions in the environmental group were selected. The resulting subpanel consists of 289 natural *Arabidopsis thaliana* accessions spanning the natural range of the species, which we dub the ‘EcoCore’ panel (**Figure 2B-C, Supplemental Dataset 6**). By selecting similar numbers of accessions from each environmental group, we ensured that each major environment within the natural distribution range of *A. thaliana* was equally represented in the EcoCore panel. This was confirmed by t-SNE analysis on the full AraClim database (**Supplemental Figure 12, Supplemental Figure 13, Supplemental Figure 14**) and the 11 non-redundant environmental parameters (**Supplemental Figure 15**), which indicated that the EcoCore panel effectively covers the variation in local environments present in the full 1001G panel. PCA analysis on the genetic map of the 1001G panel also confirmed that the EcoCore panel covers the genetic space of the 1001G panel (**Supplemental Figure 16**).

To confirm that our environment-based selection procedure indeed resulted in a more environmentally diverse panel compared to other panels, we compared the EcoCore panel to 10,000 random generated panels of the same size using the WorldClim 2 parameters (Fick & Hijmans, 2017). The EcoCore panel was not designed to maximize diversity in WorldClim 2 parameters, since this set comprises only 19 parameters (descriptors of local temperature and precipitation), compared to the 373 parameters in the AraClim dataset that were used for selecting the EcoCore panel. Nevertheless, we found that the EcoCore panel had a high SD in most of the WorldClim 2 variables compared to the 10,000 random panels (**Figure 3**) (mean percentile = 85, less than 1% of random panels had a higher mean percentile; **Supplemental figure 17**). This SD percentile was especially high for precipitation-related variables. For most of the temperature-related WorldClim 2 parameters, the EcoCore was in the top 25 percentile, except for BIO3, BIO5, BIO9 and BIO11. Performing the same analysis for the SD using all AraClim variables showed the same pattern with the EcoCore panel having a high SD rank for most, but not all, environmental variables (**Supplemental Figure 18**). Note that we did not use the SD rank as a panel selection criterium, because explicitly maximizing the SD would lead to over-representation of extreme or outlier-environments. In contrast, our clustering-based approach ensures representation across environmental regimes with realistic frequencies of conditions experienced by natural populations, while maintaining the geographical and population structure important for genotype-environment associations. Therefore, the high SD rank of the EcoCore panel is especially meaningful, because it reflects high environmental variation achieved alongside ecologically balanced sampling and preservation of geographic and population structure.

**Figure 3:**
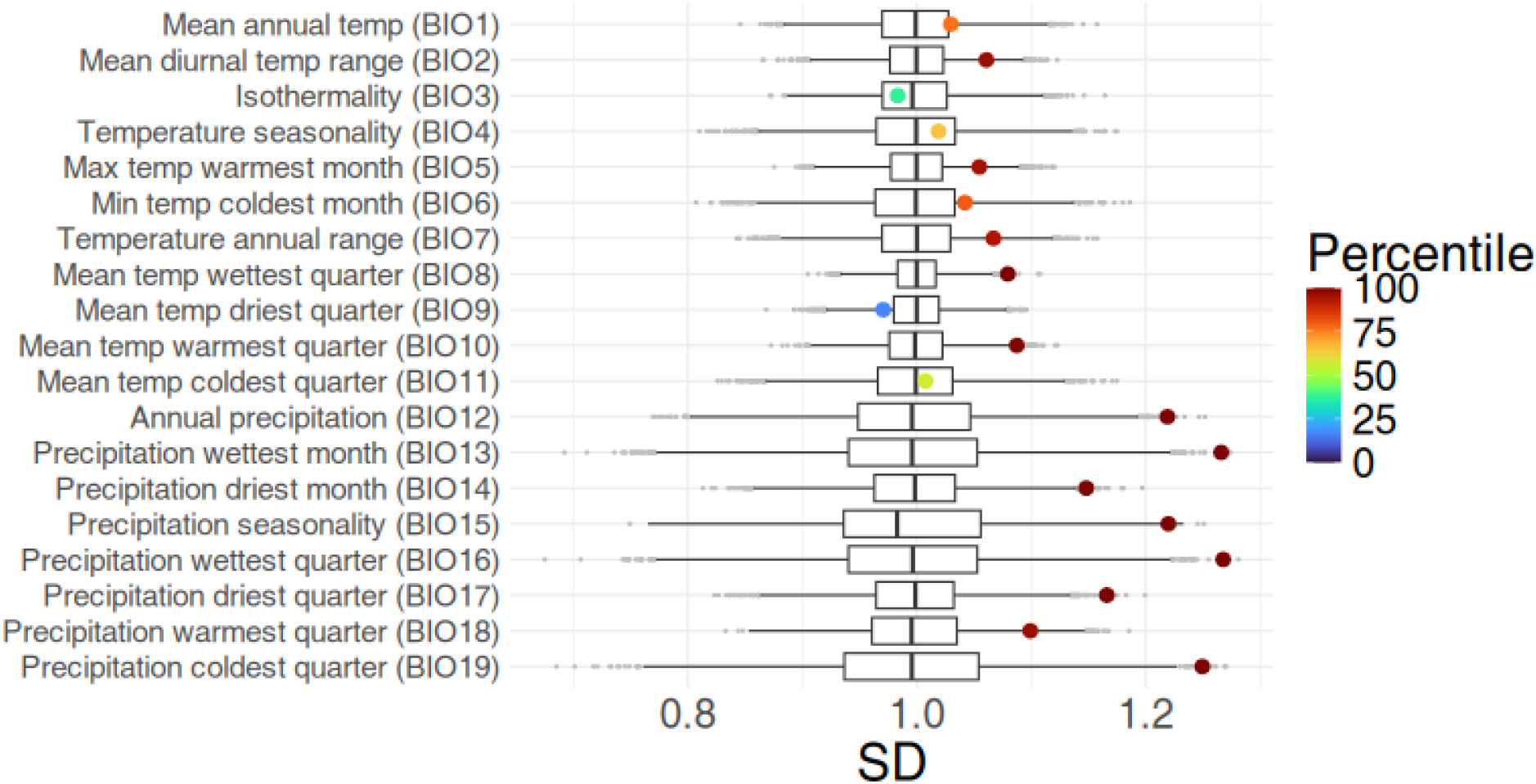
SD distribution of random subpanels and the EcoCore panel per WorldClim 2 parameter. Boxplots of the SD distributions over 10000 generated random panels of the same size as the EcoCore panel (289 accessions). Large dots indicate the SD of the EcoCore panel, with the color indicating the percentile of the EcoCore panel SD, relative to the SD of the random panels shown in the boxplots.

### Performance of EcoCore compared to the full 1001G panel

Next, we aimed to evaluate the effectiveness of our EcoCore panel in a GWAS study of environment-responsive plant traits. We focused on hypocotyl length of young seedlings in two different temperatures for several reasons. First, out of the 373 AraClim parameters temperature explained most of the environmental variation across accessions (**Supplemental Figure 4**). In addition, Arabidopsis shows signs of adaptation to the local temperature environment for several traits (Clauw et al., 2022; Hoffmann, 2002; Ibañez et al., 2017; Martínez-Berdeja et al., 2020; Wu et al., 2026). Finally, hypocotyl length is a hallmark trait of temperature-responsiveness to mild high temperatures (thermomorphogenesis), scales to temperature input of accessions, *and* is a reliable predictor of temperature responsiveness and plant performance later in life, including flowering time (Ibañez et al., 2017).

We measured hypocotyl lengths of seedlings of 966 available accessions from the 1001G panel (of which 256 overlap with the EcoCore subpanel) grown in 20°C and 28°C (**Supplemental Dataset 7, Supplemental Figure 19**). Next, we assessed how hypocotyl lengths associated to the local environment at the collection site. The 20 environmental groups that were defined to design the EcoCore panel (**Figure 2**) explained 9.6% and 11% of the variance in hypocotyl length in 20°C and 28°C respectively. We investigated if this was more than expected by chance by randomly distributing the accessions across the environmental groups 9999 times and calculating the variance explained. Indeed, the non-permutated accession sorting (i.e. when assigned to the original group) explained more variation in hypocotyl length in both temperature conditions than any of the 9999 random orderings. This indicates that observed hypocotyl lengths, even though being measured in controlled climate cabinets, highly significantly relates to the environmental conditions prevailing at the local site of collection (P = 0.0001).

Because the environmental groups used to construct the EcoCore panel were associated with hypocotyl length, we wondered whether the EcoCore would capture a larger variance in hypocotyl length than random panels of the same size. To test this, we generated 9999 random subpanels of the 1001G of the same size as the EcoCore and calculated the variance in hypocotyl length. This indicated that the EcoCore was in the 78^th^ percentile in 20°C (non-significant) the 91^st^ percentile in 28°C (P< 0.1) and in the 94^th^ percentile (P < 0.1) for the difference between the two temperature conditions (**Supplemental Figure 20**). Thus, although hypocotyl length only constitutes one out of many known environmental responses, this example shows that the EcoCore can capture high trait variance for environmental responses to conditions that are sub-optimal for most accessions (only 217 out of 1066 accessions or 5 out of 20 environments experience such high ambient temperature (28°C) according to the WorldClim 2 ‘Maximum temperature in warmest month’ parameter).

The median hypocotyl length in 20°C ranged from 2.02 mm in environmental group #14 (found in Sweden and Finland) to 2.70 mm in environmental group #16 (found in The Netherlands, Germany, Kirgizstan) (**Supplemental Figure 21A**), while in 28°C the median hypocotyl length differed between 4.92 mm in environment #14 and 7.36 in environment #3 (found in the east coast of north America, and central Europe) (**Supplemental Figure 21B**). As expected, we found a strong correlation between the hypocotyl length in 20°C and the hypocotyl length in 28°C (Pearson correlation = 0.75), but with some variation in the response to increased temperature (**Supplemental figure 21C**). The EcoCore panel roughly covered the phenotypic range of the full 1001G panel (**Supplemental figure 21D**). The strongest correlation between hypocotyl length and any individual environmental parameter (of the 373 AraClim2 parameters assessed) was observed between hypocotyl length in 20°C and predicted evapotranspiration (water loss from the plant to the atmosphere; Pearson correlation = 0.26) (**Supplemental figure 21E**).

### EcoCore panel reveals Quantitative Trait Loci in environmental-response traits

Next, we assessed the effectiveness of the EcoCore panel in Quantitative Trait Loci (QTL) detection compared to the full 1001G panel. To this aim, GWAS was performed on the hypocotyl length obtained in 20°C using data of the full 1001G panel (966 accessions) (**Figure 4A**) and the EcoCore subpanel (256 accessions) (**Figure 4B**). Using the 1001G panel we found 117 significant genetic variants (–log10(P) > 6) distributed over 14 distinct peaks, with a maximum of -log10(P) of 7.55, while using the EcoCore subpanel we found 17 genetic variants distributed over 10 peaks with a maximum -log10(P) of 7.86. Interestingly, none of the SNPs nor peaks were overlapping between the two panels and each panel revealed unique genetic variation associated with hypocotyl length in 20°C. In 28°C we detected 26 significant variants distributed over 13 peaks with a maximum -log10(P) of 7.13 using the full 1001G panel and 16 significant variants over 10 peaks with a maximum -log10(P) of 7.09 using the EcoCore panel (**Supplemental Figure 22**). Only one genetic variant at chromosome 1 (0.585122 Mbp), where the alternative allele was associated with increased hypocotyl length in 28°C (mean reference = 6.3 mm, mean alternative = 7.0 mm) (**Supplemental Figure 23A**) was detected using both panels. 9.4% of accessions of the 1001G from 12 different environmental groups had the alternative allele (**Supplemental Figure 24A**), while in the EcoCore panel 10.2% of accessions from 11 different environmental groups had the alternative allele (**Supplemental Figure 24B**). The alternative allele was rare or absent in most environmental groups, but common in environmental group #13 (47 out of 66 of accessions, 71.2%) (**Supplemental Figure 24**), particularly in accessions from northern Sweden (**Supplemental Figure 23B**). This variant was located inside the gene *IMPORTIN ALPHA ISOFORM 6 (IMPα-6;* At1G02690). *IMPα-6* promotes the nuclear transport of *MYB DOMAIN PROTEIN 3R-4* (MYB3R4) (Yang et al., 2021), and *myb3r4* mutants exhibit very short hypocotyls (Haga et al., 2011). Furthermore, in rice *OsMYB3R-2* has been linked to temperature adaptation (albeit cold stress) (Ma et al., 2009), suggesting a possible effect of *IMPα-6* on hypocotyl length in 28°C.

The GWAS results revealed an EcoCore-specific peak with high significance (–log10(P) = 7.86) on chromosome 5 at ∼20.4 Mbp for hypocotyl length in 20°C (**Figure 4A**, see **Supplemental Figure 25**) for zoom in-on locus), whereof the alternative allele is associated with longer hypocotyls (**Supplemental Figure 26**). To assess why this peak was identified with the EcoCore panel but not the 1001G panel, we compared the geographic distribution of the environmental groups (**Figure 5A**) with the distribution of the alternative allele (**Figure 5B**). Notably, 12 out of 27 accessions with the alternative allele belong to environmental group #9 (exclusively in the UK), or environmental group #15 (UK and north-western Europe) (**Figure 5A-B**), although being present in several groups at low frequencies (**Figure 5C**). Out of these, environmental group #15 was relatively small, containing only 31 accessions (3.4% of the 1001G). Thus, by design, taking an equal number of accessions from the 20 diverse environmental groups **(Figure 5B)**, the EcoCore enriched accessions from ‘rare’ environmental groups such as group #15. As a consequence, also alleles occurring in these small environmental groups become enriched, as is the case for this GWAS peak wherein 2.8% of accessions had the alternative allele in the 1001G panel versus 5.1% in the EcoCore panel (**Figure 5D**).

**Figure 4:**
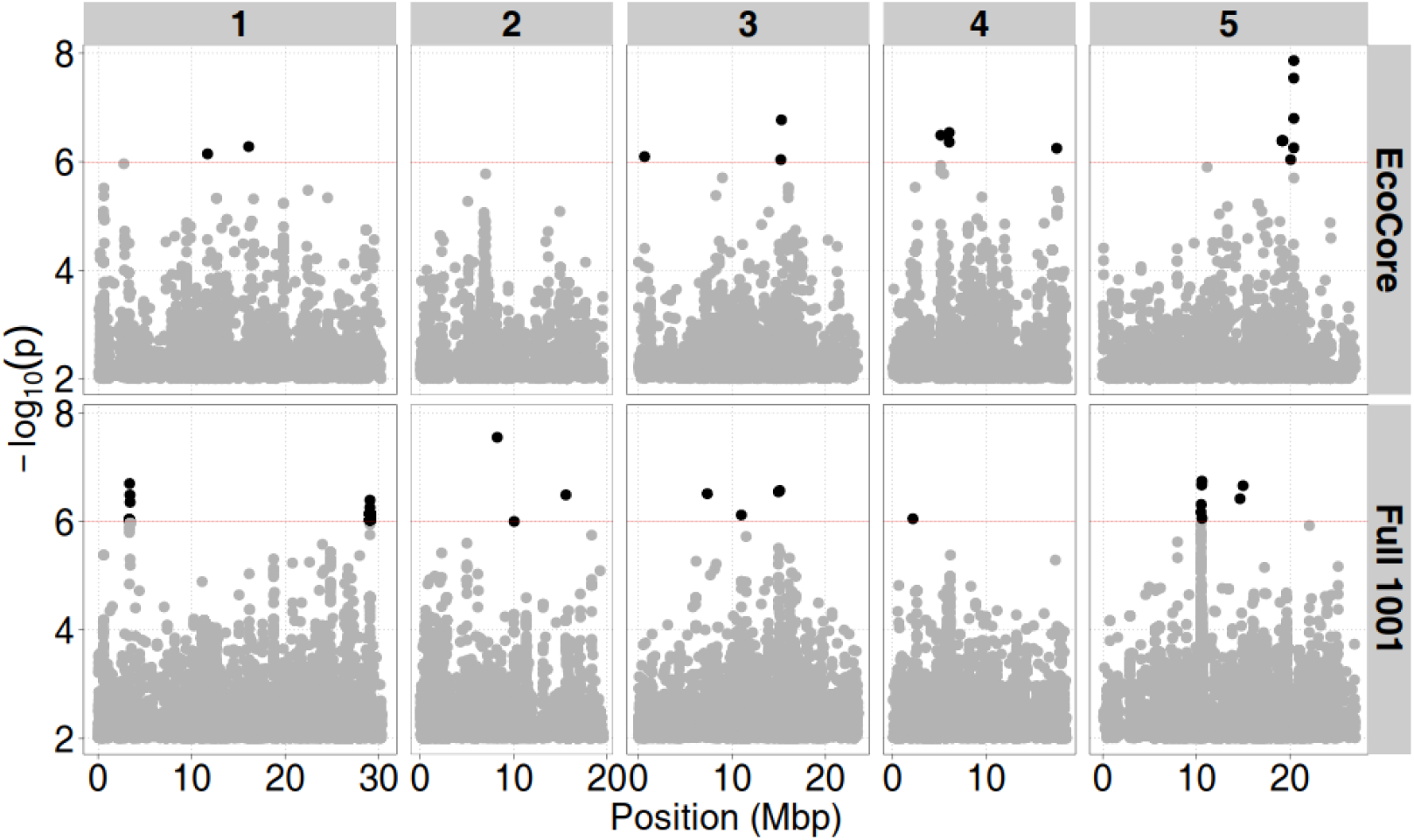
GWAS results for hypocotyl length in 20°C. Indicated are associations mapped with **A)** the 1001G panel (966 accessions) and **B)** mapped with the EcoCore panel (256 accessions). -log_10_(p) indicates the significance of the mapped genetic variants across the 5 chromosomes of Arabidopsis (genetic position indicated in mega bases pairs; Mbp). The red line indicates the threshold of significance, above which significant associations are indicated in bold.

**Figure 5:**
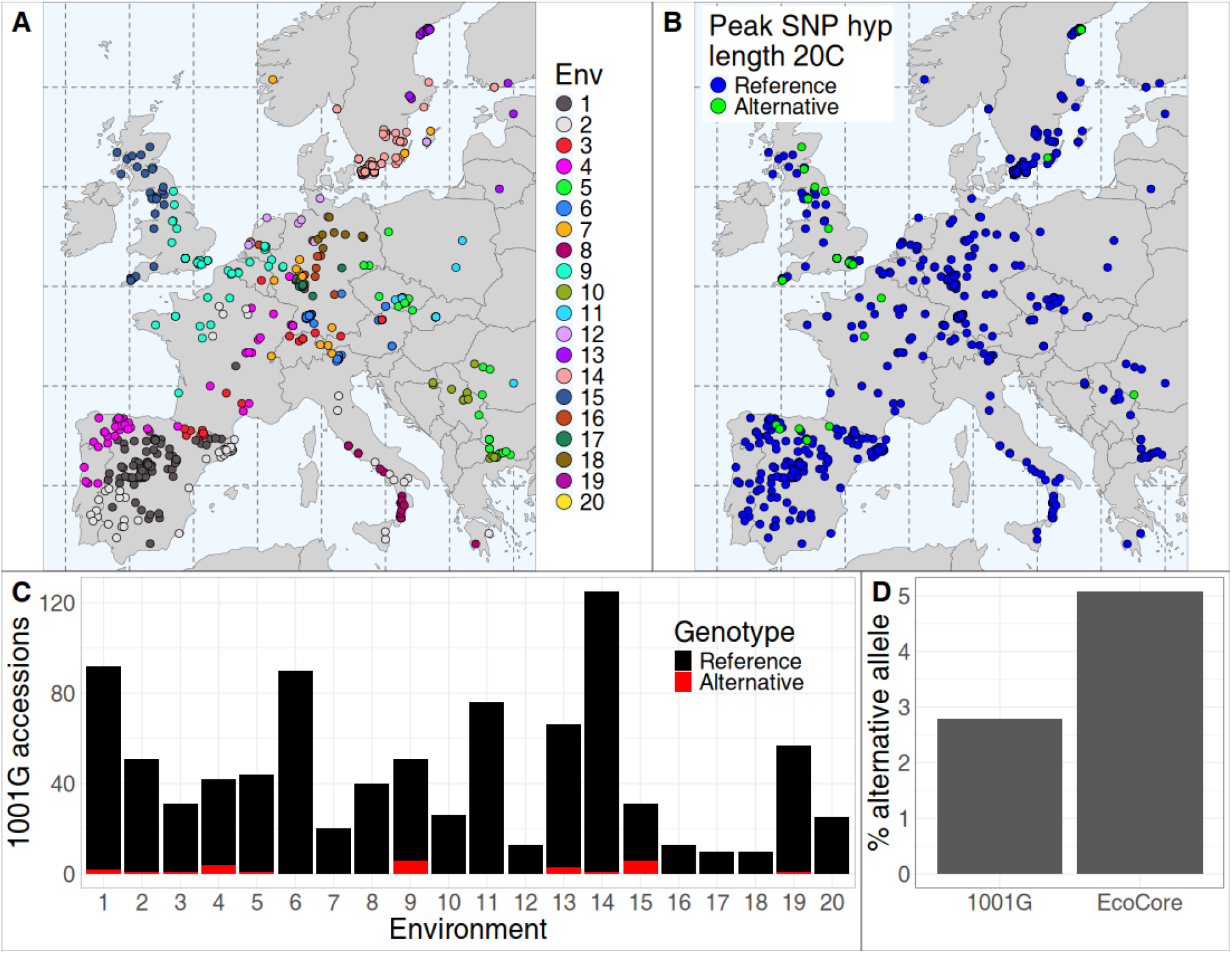
The EcoCore panel enriches a rare allele associated with hypocotyl length at 20°C. **A, B)** Maps showing all European 1001G accessions. Colors indicate **A)** environmental group and **B)** allele variant of the peak SNP for hypocotyl length in 20°C on chromosome 5; 20.429829 Mbp, identified using the EcoCore panel, but not the 1001G panel (blue indicates reference allele, green indicates alternative allele). **C)** Allele distribution per environmental group per allele (black indicates reference allele, red indicates alternative allele). **D)** frequency of accessions with the alternative allele in the 1001G and the EcoCore panels.

By enriching accessions from smaller environmental groups, the EcoCore panel also enriches trait values occurring in these smaller environmental groups. In our hypocotyl length data this is not so apparent, since both large and small median hypocotyl lengths are found in large environmental groups. For example, the smallest median hypocotyl length is found in the largest environmental group (#14, predominantly Swedish accessions) whereas the second longest median hypocotyl length is found in the second largest environmental group (#1, predominantly Iberian accessions) **(Supplemental figure 21)**. To further confirm that the EcoCore can enrich trait values from rare environments, we assessed existing data on flowering time in 10°C and 16°C (scored for 943 and 916 accessions of the 1001G and 276 and 271 accessions of the EcoCore respectively (The 1001 Genomes Consortium et al., 2016; Zan & Carlborg, 2019), obtained from the AraPheno database. Flowering time depends on local environmental conditions and extensive (adaptive) natural variation exists (Fournier‐Level et al., 2022; Kang et al., 2023; Zan & Carlborg, 2019). These traits indeed depended on the local environment (**Supplemental figure 27A, Supplemental figure 27B**). Strikingly, the lowest values occur in the three smallest environmental groups (#16, #17, and #18), creating a distribution skewed towards faster flowering times in the EcoCore panel compared to the 1001G panel (**Supplemental figure 27C, Supplemental figure 27D**). Hence, the EcoCore can indeed enrich for environmental trait values that have few representative accessions in the 1001G, facilitating the discovery of novel associations that help plants cope with specific or understudied environments and that are not likely to be detected using the full 1001G panel.

## Discussion

(Sub)panels of *Arabidopsis thaliana* accessions are usually selected from a larger population based on maximization of genetic variation (Li et al., 2010). While this is intuitive, alleles need to reach a sufficient frequency for quantitative genetic assessment (e.g. GWAS) to be efficient (commonly considered by filtering SNPs based on minor allele frequency (Floriani & Lipka, 2025). Here, we have shown that the number of such alleles is an asymptotic function of the number of accessions within a given set, with only a minor effect exerted by the specific panel composition (**Figure 1B**). Our results indicate that instead of maximizing genetic dissimilarity, it is at least equally valid to consider other attributes of the accessions as design principle for a subset. Building on this idea, we designed a panel based on optimizing the variation in the environment, which we named the EcoCore panel. We showed that the EcoCore panel captured more environmental variance of the WorldClim 2 variables and more trait variance for hypocotyl lengths in 20°C and 28°C than random sets derived from the full 1001G panel, and led to novel hypocotyl elongation QTLs. We hence put forward the EcoCore panel as a resource for studying the genetics of environmental responses. We deem that a standardized panel for studying environmental responses will enhance the consensus, reproducibility and integration of results, while at the same time recognizing that studying a specific environmental response may be best done by selecting accessions based on that specific environmental variable.

Variation in environmental parameters may correlate with geographic and genetic variation (Lasky et al., 2012). Nevertheless, selecting accessions from diverse environments is not the same as selecting accessions from different geographic regions or genetic groups, because 1) geographically distant areas can experience very similar climates whereas (micro)environments can change on a small geographic scale and 2) because similar environments can lead to very fast convergent evolution (Wu et al., 2026). For instance, some European accessions experience similar conditions to North American accessions (**Supplemental Figure 10**). However, genetics, adaptations, and environmental (dis)similarity cannot be seen as entirely independent entities either, as accessions that are geographically close (and thus persist in relatively similar environments) are usually also genetically relatively similar because of shared ancestry (isolation by distance) (Platt et al., 2010). Overall, selecting a subpanel based on environmental variation, as we did for the EcoCore, is different from most existing approaches that are based on solely genetic criteria and may enrich for adaptive alleles. An alternative strategy that has been effective in identifying variants associated with environmental responses is to focus on specific geographic subpopulations that range over contrasting environments. For example, (Satbhai et al., 2017) used Swedish 1001 Genomes panel accessions, which range over contrasting soil conditions, to disclose the genetics underlying iron limitation response. Likewise, the ‘Dutch 1001G’ accessions have been used to uncover variants and genes linked to wind resistance, iron deficiency tolerance, and drought tolerance (Wijfjes et al., 2024). Focusing on specific populations reduces confounding effects from population structure and complex genetic interactions, but its potential for discovery is limited because sampling a narrow geographic subpopulation is prone to capturing far less natural and genetic variation than a panel spanning the (close-to) full range of *A. thaliana* geographic distribution.

We used the full 1001G panel and the EcoCore panel to study the genetics of hypocotyl elongation in different temperature regimes (20°C and 28°C). We noticed that both panels yielded mainly unique QTLs due to different allele distributions between the panels (**Figure 4, Supplemental Figure 28**). This included a QTL for hypocotyl length in 20°C found only with the EcoCore panel. In general, any set of accessions will cause certain alleles to be underrepresented and others to be overrepresented relative to the entire population. This confounding effect will lead to novel or missed associations per definition. Even different panels of the same size might lead to very different associations due to different allele distributions and interactions between them, the overall population structure, and complex phenotypes. In that sense, a sub-panel always represents a trade-off and the optimal subpanel theoretically does not exist. Even the 1001G panel is a biased subpanel of the total *Arabidopsis thaliana* population in the wild and therefore subject to these same trade-offs. However, we showed that the peak SNP associated with hypocotyl length in 20°C occurred mostly in rare environments (**Figure 5**), and such alleles can be enriched in the EcoCore because it was designed based on equal representation of environments. This balances for a particular confounding effect of the 1001G, which is biased towards the Swedish and Iberian populations (**Supplemental Figure 11**). These are also seemingly the populations which have been most studied in isolation and their genetics has been most extensively explored (Brachi et al., 2025; Castilla et al., 2020; Huber et al., 2014; Kerdaffrec et al., 2016; Kerdaffrec & Nordborg, 2017; Méndez-Vigo et al., 2011; Méndez‐Vigo et al., 2013; Picó et al., 2008; Subrahmaniam et al., 2025; Toledo et al., 2020). Therefore, by enriching accessions from rare environments, the EcoCore panel also enriches for genetic variation underrepresented in the 1001G, enhancing the chance of discovering novel genes driving environmental responsiveness.

Surprisingly, the strongest correlation between hypocotyl length and any individual environmental parameter (out of the 373 AraClim2 parameters assessed) was between the hypocotyl length in 20°C and predicted evapotranspiration (water loss from the plant to the atmosphere) (Pearson correlation = 0.26) **(Supplemental figure 21E)**, suggesting short hypocotyls may attribute to water conservation, or that short hypocotyls scale with compact plants, leading to water conservation. The latter is indirectly supported by the notion that the opposite phenotype, being long hypocotyls, is a predictor of a thermomorphogenic ‘open’ plant architecture (Ibañez et al., 2017), and thermomorphogenesis attributes to improved plant evaporative cooling under warm temperatures (Crawford et al., 2012; Park et al., 2019). However, when one considers functional explanations like this, or more generally aims at connecting environment to location of origin, one should realize that the geographic locations of the reported accessions are not in all cases very precise. In addition, maps of environmental parameters are made by interpolating between weather stations and therefore accuracy depends on the density of weather stations and the scale of local climate variation. Furthermore, geospatial data is obtained with a certain spatial resolution that may prevent capturing the exact (micro)environment at the collection site. Put in other words, the environmental factors that are associated with each accession come with a healthy degree of uncertainty.

In summary, we showed that in subpanels of the 1001G panel the composition does not substantially affect the number of varying alleles, and this finding motivated us to instead select a panel based on equal representation from different collection environments. We measured hypocotyl length in 20°C and 28°C for almost the full 1001G panel and showed that the EcoCore panel captures more trait variance compared to random sub-panels of the 1001G, which aids GWAS on traits affected by the environment. Overall, this study both enhances understanding of the genetic basis of hypocotyl length in different temperature regimes and presents the EcoCore as a resource for studying environmental response traits.

## Materials and methods

### Code and figures

All analyses were done in R 4.3.1 (R Core Team, 2022). Figures were made using ggplot2 (Martin et al., 2022) and combined using the cowplot package (Claus O. Wilke, 2025). Maps showing the geographic location of *Arabidopsis* accessions were made using the naturalearth package (Massicotte & South, 2017). We used the tidyr package for coding convenience and the parallel package to distribute computing tasks over multiple cores (R Core Team, 2025). Correlation networks were made using code from (van Eijnatten et al., 2025) based on the igraph, ggraph, and tidygraph packages (Csardi & Nepusz, 2006; Thomas Lin Pedersen, 2024b, 2024a). Heatmaps were colored using the “vik” palette from the scico package (Pedersen & Crameri, 2018).

### Investigating typical panel sizes

Phenotypic data for the 1001G accessions (435 phenotypes from 25 studies) was obtained from the AraPheno database (Seren et al., 2017; Togninalli et al., 2019) and a curated extension of the AraPheno database (1864 phenotypes) was obtained from (Ruffley et al., 2023). We removed climate (not measured) phenotypes from the datasets. Both datasets were used to investigate panel sizes per study. We noticed that the panel size in AraPheno was often lower than reported in the source literature because many panels included accessions not part of the 1001G panel. Hence, we gathered the panel size of the 25 studies in AraPheno from source literature and reported this in figure 1A.

### Available genetic variation of random panels

We obtained the SNP matrix of the full 1001 panel from (Arouisse et al., 2020). For each possible panel size between 100 and 1066 accessions we generated 100 random panels (for a total of 96700 random panels) and calculated the number of genetic variants where at least 13 accessions had the reference allele and at least 13 accessions had the alternative allele.

### Collecting environmental data

Local environment data and geographic coordinates were obtained from the AraClim v2 database (Ferrero-Serrano & Assmann, 2019). This database contains measured and predicted climate and environment variables from various sources (satellites, models, etc.). Most variables concern temperature, precipitation or soil characteristics. In addition, WorldClim 2 (Fick & Hijmans, 2017) raster data was downloaded using the worlclim_global() function from the geodata package (Robert J. Hijmans, 2025) (at 0.5 degree-minute resolution) and extracted from the raster object using the raster- and sp-R packages (Edzer J. Pebesma and Roger Bivand, 2005; Robert J. Hijmans, 2024). Accessions with no WorldClim 2 data due to missing or faulty coordinates (e.g. in the ocean) and AraClim v2 variables with more than 10 missing values were not included in analysis (leaving 1066 accessions and 373 environmental variables).

### Selecting non-redundant environmental variables

Principal component analysis on the scaled AraClim v2 matrix was performed using the prcomp() function from the stats package (**Supplemental Figure 4**). To reduce the dimensionality of the data without losing interpretability, we clustered the AraClim v2 variables by performing k-means with the kmeans() function from base R on the AraClim v2 correlation matrix and selected for each cluster the environmental variable closest to the cluster centroid to get 11 representative environmental variables.

### EcoCore panel construction

The accessions in the 1001G panel were correlated based on the 11 scaled environmental variables and hierarchically-clustered on the correlation matrix using the hclust() function from the stats package. 20 environmental groups of accessions were defined from the dendrogram using the cutree() function with k = 20. To generate the EcoCore set, we selected 15 accessions per environmental cluster by minimizing the genetic covariance between accessions. If there were less than 15 accessions in an environment, we selected all. Genetic covariance was maximized by calculating the covariance matrix of the SNP matrix (Arouisse et al., 2020) using the cov() function from the stats function and performing a hierarchical clustering on the covariance matrix using the hclust() function. 15 accessions were then chosen by defining 15 genetic groups from the dendrogram using the cutree() function with k = 15 and one accession was selected randomly from each genetic group. This procedure resulted in a total of 289 accessions included in the EcoCore panel.

### Hypocotyl elongation essays

Seeds of the 1001 genomes set were obtained via the European Arabidopsis Stock Centre (NASC). After one round of propagation in long day conditions (16h light and 8h darkness) at 20°C, in some cases preceded by a vernalization period (short day; 8h light and 16h darkness, 2°C-7°C) of 4 weeks, the seeds were surface sterilized with 70% ethanol for 4 min and with 4% sodium hypochlorite solution for 8 min. Subsequently, the seeds were stratified at 4°C in water for 4 days. Subsequently, the sterile seeds were sown on *Arabidopsis thaliana* salts (ATS) medium (Lincoln et al., 1990) in the presence of 1% sucrose on Petri plates. Per accession two replicate ATS plates were prepared and each plate contained 2 rows of 20 seeds. Subsequently, the plates were cultivated vertically for 4 days at 20 °C, under 16 h long day photoperiod at 90 µMol m^2^ sec^-1^ light (fluorescent tubes, Osram HO 39W/840 LUMILUX Cool White) growth cabinet (Conviron). After these 4 days, late-/non-germinators were marked and one of the plates per accession were shifted to a growth cabinet set at 28°C while the other plate remained at 20°C for 4 additional days. After a total of 8 days the plates were photographed using a Canon EOS 700D Camera and the Canon EOS Utility software, and hypocotyl lengths were measured using the RootDetection 0.1.3-beta-2 software (https://labutils.de/rd_download.html). In total, 983 individual genotypes were sown, of which 966 could be evaluated. Of these 966, 256 accessions were part of the EcoCore set, thus covering in total 89% of 289 EcoCore accessions. We filtered the hypocotyl lengths for seedlings that deviated more than 2x SD from the accession mean. We calculated for each accession the mean for the filtered hypocotyls and used this as traits for GWAS.

### GWAS

The GWAS approach was based on the lme4QTL package (Ziyatdinov et al., 2018) as described in (Dijkhuizen et al., 2025). Population structure corrections were done by using the covariance matrix of the SNP matrix (obtained with the cov() function) and by adding the first five principal components of the SNP matrix (obtained with prcomp() function) as cofactor in the GWAS. When mapping hypocotyl length we used a MAF threshold of 0.05 for the EcoCore panel resulting in 1792373 SNPs for 256 accessions in the EcoCore panel. When mapping with the full 1001G panel (966 accessions) we used a MAF of 0.27 (at least 25 of each allele) to not miss rare alleles, which lead to 1785441 SNPs.

## Author contributions

ALvE, JJK, MvZ and LBS conceived the project, ALvE, JJK, MvZ and LBS analyzed data, ALvE and LBS visualized the data, JT, CD, MQ performed and analyzed hypocotyl length assays, MvZ and LBS supervised the project. ALvE, JJK, MvZ and LBS wrote the paper, with input of all authors.

## Acknowledgements

The authors like to thank colleagues of the CropXR program for their valuable insights and sharing their thoughts during early stages of the EcoCore panel development. The authors also thank Kathrin Denk for her substantial contribution in the lab during the hypocotyl length assays.

## Data and Code availability

The Supplementary Datasets including gathered phenotypes and environmental parameters as well as the measured hypocotyl lengths can be found in Supplementary_Datasets.xlsx.

## References

Ågren, J., & Schemske, D. W. (2012). Reciprocal transplants demonstrate strong adaptive differentiation of the model organism Arabidopsis thaliana in its native range. New Phytologist, 194(4), 1112–1122. 10.1111/j.1469-8137.2012.04112.x

Arouisse, B., Korte, A., van Eeuwijk, F., & Kruijer, W. (2020). Imputation of 3 million SNPs in the Arabidopsis regional mapping population. The Plant Journal, 102(4), 872–882. 10.1111/tpj.14659

Brachi, B., Filiault, D. L., Pisupati, R., Dahan-Meir, T., Igolkina, A., Anastasio, A., Box, M. S., Duncan, S., Karasov, T. L., Kerdaffrec, E., Merwin, L., Morton, T. C., Nizhynska, V., Novikova, P. Yu., Rabanal, F., Tsuchimatsu, T., Säll, T., Dean, C., Holm, S., … Nordborg, M. (2025). Life-history trade-offs explain local adaptation in Arabidopsis thaliana. 10.1101/2025.05.18.654693

Brady, S., Auge, G., Ayalew, M., Balasubramanian, S., Hamann, T., Inze, D., Saito, K., Brychkova, G., Berardini, T. Z., Friesner, J., Ho, C., Hauser, M., Kobayashi, M., Lepiniec, L., Mähönen, A. P., Mutwil, M., May, S., Parry, G., Rigas, S., … Provart, N. J. (2025). Arabidopsis research in 2030: Translating the computable plant. The Plant Journal, 121(5). 10.1111/tpj.70047

Castilla, A. R., Méndez-Vigo, B., Marcer, A., Martínez-Minaya, J., Conesa, D., Picó, F. X., & Alonso-Blanco, C. (2020). Ecological, genetic and evolutionary drivers of regional genetic differentiation in Arabidopsis thaliana. BMC Evolutionary Biology, 20(1), 71. 10.1186/s12862-020-01635-2

Claus O. Wilke. (2025). cowplot: Streamlined Plot Theme and Plot Annotations for “ggplot2.”

Clauw, P., Coppens, F., De Beuf, K., Dhondt, S., Van Daele, T., Maleux, K., Storme, V., Clement, L., Gonzalez, N., & Inzé, D. (2015). Leaf Responses to Mild Drought Stress in Natural Variants of Arabidopsis . Plant Physiology, 167(3), 800–816. 10.1104/pp.114.254284

Clauw, P., Kerdaffrec, E., Gunis, J., Reichardt-Gomez, I., Nizhynska, V., Koemeda, S., Jez, J., & Nordborg, M. (2022). Locally adaptive temperature response of vegetative growth in Arabidopsis thaliana. ELife, 11. 10.7554/eLife.77913

Crawford, A. J., McLachlan, D. H., Hetherington, A. M., & Franklin, K. A. (2012). High temperature exposure increases plant cooling capacity. Current Biology, 22(10), R396–R397. 10.1016/j.cub.2012.03.044

Csardi, G., & Nepusz, T. (2006). The igraph software package for complex network research. InterJournal Complex Systems, Complex Sy(1695).

Dijkhuizen, R., van Eijnatten, A. L., Mehrem, S. L., van den Bergh, E., van Lieshout, J., Spaninks, K., Kaandorp, S., Offringa, R., Proveniers, M., van den Ackerveken, G., & Snoek, B. L. (2025). From aerial drone to quantitative trait locus: leveraging next-generation phenotyping to reveal the genetics of color and height in field-grown Lactuca sativa. The Plant Journal, 123(3). 10.1111/tpj.70405

Donohue, K., Heschel, M. S., Butler, C. M., Barua, D., Sharrock, R. A., Whitelam, G. C., & Chiang, G. C. K. (2008). Diversification of phytochrome contributions to germination as a function of seed-maturation environment. New Phytologist, 177(2), 367–379. 10.1111/j.1469-8137.2007.02281.x

Edzer J. Pebesma and Roger Bivand. (2005). Classes and methods for spatial data in R.

Exposito-Alonso, M., Burbano, H. A., Bossdorf, O., Nielsen, R., & Weigel, D. (2019). Natural selection on the Arabidopsis thaliana genome in present and future climates. Nature, 573(7772), 126–129. 10.1038/s41586-019-1520-9

Ferrero-Serrano, Á., & Assmann, S. M. (2019). Phenotypic and genome-wide association with the local environment of Arabidopsis. Nature Ecology & Evolution, 3(2), 274–285. 10.1038/s41559-018-0754-5

Fick, S. E., & Hijmans, R. J. (2017). WorldClim 2: new 1-km spatial resolution climate surfaces for global land areas. International Journal of Climatology, 37(12), 4302–4315. 10.1002/joc.5086

Floriani, T., & Lipka, A. E. (2025). Rare variants in crops: theoretical insights and emerging detection strategies. In Silico Plants, 7(2). 10.1093/insilicoplants/diaf012

Fournier-Level, A., Taylor, M. A., Paril, J. F., Martínez-Berdeja, A., Stitzer, M. C., Cooper, M. D., Roe, J. L., Wilczek, A. M., & Schmitt, J. (2022). Adaptive significance of flowering time variation across natural seasonal environments in Arabidopsis thaliana. New Phytologist, 234(2), 719–734. 10.1111/nph.17999

Haga, N., Kobayashi, K., Suzuki, T., Maeo, K., Kubo, M., Ohtani, M., Mitsuda, N., Demura, T., Nakamura, K., Jürgens, G., & Ito, M. (2011). Mutations in MYB3R1 and MYB3R4 Cause Pleiotropic Developmental Defects and Preferential Down-Regulation of Multiple G2/M-Specific Genes in Arabidopsis. Plant Physiology, 157(2), 706–717. 10.1104/pp.111.180836

Hoffmann, M. H. (2002). Biogeography of Arabidopsis thaliana (L.) Heynh. (Brassicaceae). Journal of Biogeography, 29(1), 125–134. 10.1046/j.1365-2699.2002.00647.x

Hopkins, R., Schmitt, J., & Stinchcombe, J. R. (2008). A latitudinal cline and response to vernalization in leaf angle and morphology in Arabidopsis thaliana (Brassicaceae). New Phytologist, 179(1), 155–164. 10.1111/j.1469-8137.2008.02447.x

Huber, C. D., Nordborg, M., Hermisson, J., & Hellmann, I. (2014). Keeping It Local: Evidence for Positive Selection in Swedish Arabidopsis thaliana. Molecular Biology and Evolution, 31(11), 3026–3039. 10.1093/molbev/msu247

Ibañez, C., Poeschl, Y., Peterson, T., Bellstädt, J., Denk, K., Gogol-Döring, A., Quint, M., & Delker, C. (2017). Ambient temperature and genotype differentially affect developmental and phenotypic plasticity in Arabidopsis thaliana. BMC Plant Biology, 17(1), 114. 10.1186/s12870-017-1068-5

Jiang, Z., van Zanten, M., & Sasidharan, R. (2025). Mechanisms of plant acclimation to multiple abiotic stresses. Communications Biology, 8(1), 655. 10.1038/s42003-025-08077-w

Jiang, Z., Verhoeven, A., Li, Y., Geertsma, R., Sasidharan, R., & van Zanten, M. (2024). Deciphering acclimation to sublethal combined and sequential abiotic stresses in Arabidopsis thaliana. Plant Physiology. 10.1093/plphys/kiae581

Johanson, U., West, J., Lister, C., Michaels, S., Amasino, R., & Dean, C. (2000). Molecular Analysis of FRIGIDA , a Major Determinant of Natural Variation in Arabidopsis Flowering Time. Science, 290(5490), 344–347. 10.1126/science.290.5490.344

Kang, M., Wu, H., Liu, H., Liu, W., Zhu, M., Han, Y., Liu, W., Chen, C., Song, Y., Tan, L., Yin, K., Zhao, Y., Yan, Z., Lou, S., Zan, Y., & Liu, J. (2023). The pan-genome and local adaptation of Arabidopsis thaliana. Nature Communications, 14(1), 6259. 10.1038/s41467-023-42029-4

Kerdaffrec, E., Filiault, D. L., Korte, A., Sasaki, E., Nizhynska, V., Seren, Ü., & Nordborg, M. (2016). Multiple alleles at a single locus control seed dormancy in Swedish Arabidopsis. ELife, 5. 10.7554/eLife.22502

Kerdaffrec, E., & Nordborg, M. (2017). The maternal environment interacts with genetic variation in regulating seed dormancy in Swedish Arabidopsis thaliana. PLOS ONE, 12(12), e0190242. 10.1371/journal.pone.0190242

Koornneef, M., Alonso-Blanco, C., & Vreugdenhil, D. (2004). Naturally occurring genetic variation in Arabidopsis thaliana. Annual Review of Plant Biology, 55(1), 141–172. 10.1146/annurev.arplant.55.031903.141605

Lasky, J. R., Des Marais, D. L., McKay, J. K., Richards, J. H., Juenger, T. E., & Keitt, T. H. (2012). Characterizing genomic variation of Arabidopsis thaliana: the roles of geography and climate. Molecular Ecology, 21(22), 5512–5529. 10.1111/j.1365-294X.2012.05709.x

Lee, I., Michaels, S. D., Masshardt, A. S., & Amasino, R. M. (1994). The late-flowering phenotype of FRIGIDA and mutations in LUMINIDEPENDENS is suppressed in the Landsberg erecta strain of Arabidopsis. The Plant Journal, 6(6), 903–909. 10.1046/j.1365-313X.1994.6060903.x

Li, Y., Huang, Y., Bergelson, J., Nordborg, M., & Borevitz, J. O. (2010). Association mapping of local climate-sensitive quantitative trait loci in Arabidopsis thaliana. Proceedings of the National Academy of Sciences, 107(49), 21199–21204. 10.1073/pnas.1007431107

Ma, Q., Dai, X., Xu, Y., Guo, J., Liu, Y., Chen, N., Xiao, J., Zhang, D., Xu, Z., Zhang, X., & Chong, K. (2009). Enhanced Tolerance to Chilling Stress in OsMYB3R-2 Transgenic Rice Is Mediated by Alteration in Cell Cycle and Ectopic Expression of Stress Genes. Plant Physiology, 150(1), 244–256. 10.1104/pp.108.133454

Martin, V., Schbath, S., & Hennequet-Antier, C. (2022). R graphics with ggplot2.

Martínez-Berdeja, A., Stitzer, M. C., Taylor, M. A., Okada, M., Ezcurra, E., Runcie, D. E., & Schmitt, J. (2020). Functional variants of DOG1 control seed chilling responses and variation in seasonal life-history strategies in Arabidopsis thaliana. Proceedings of the National Academy of Sciences, 117(5), 2526–2534. 10.1073/pnas.1912451117

Massicotte, P., & South, A. (2017). rnaturalearth: World Map Data from Natural Earth. In CRAN: Contributed Packages. 10.32614/CRAN.package.rnaturalearth

McKhann, H. I., Camilleri, C., Bérard, A., Bataillon, T., David, J. L., Reboud, X., Le Corre, V., Caloustian, C., Gut, I. G., & Brunel, D. (2004). Nested core collections maximizing genetic diversity in Arabidopsis thaliana. The Plant Journal, 38(1), 193–202. 10.1111/j.1365-313X.2004.02034.x

Méndez-Vigo, B., Gomaa, N. H., Alonso-Blanco, C., & Xavier Picó, F. (2013). Among- and within-population variation in flowering time of Iberian Arabidopsis thaliana estimated in field and glasshouse conditions. New Phytologist, 197(4), 1332–1343. 10.1111/nph.12082

Méndez-Vigo, B., Picó, F. X., Ramiro, M., Martínez-Zapater, J. M., & Alonso-Blanco, C. (2011). Altitudinal and Climatic Adaptation Is Mediated by Flowering Traits and FRI, FLC, and PHYC Genes in Arabidopsis. Plant Physiology, 157(4), 1942–1955. 10.1104/pp.111.183426

Montesinos-Navarro, A., Wig, J., Xavier Pico, F., & Tonsor, S. J. (2011). Arabidopsis thaliana populations show clinal variation in a climatic gradient associated with altitude. New Phytologist, 189(1), 282– 294. 10.1111/j.1469-8137.2010.03479.x

Morales, A., de Boer, H. J., Douma, J. C., Elsen, S., Engels, S., Glimmerveen, T., Sajeev, N., Huber, M., Luimes, M., Luitjens, E., Raatjes, K., Hsieh, C., Teapal, J., Wildenbeest, T., Jiang, Z., Pareek, A., Singla-Pareek, S., Yin, X., Evers, J., … Sasidharan, R. (2022). Effects of sublethal single, simultaneous and sequential abiotic stresses on phenotypic traits of Arabidopsis thaliana. AoB PLANTS, 14(4). 10.1093/aobpla/plac029

Park, Y.-J., Lee, H.-J., Gil, K.-E., Kim, J. Y., Lee, J.-H., Lee, H., Cho, H.-T., Vu, L. D., De Smet, I., & Park, C.-M. (2019). Developmental Programming of Thermonastic Leaf Movement. Plant Physiology, 180(2), 1185–1197. 10.1104/pp.19.00139

Pedersen, T. L., & Crameri, F. (2018). scico: Colour Palettes Based on the Scientific Colour-Maps. In CRAN: Contributed Packages. 10.32614/CRAN.package.scico

Picó, F. X., Méndez-Vigo, B., Martínez-Zapater, J. M., & Alonso-Blanco, C. (2008). Natural Genetic Variation of Arabidopsis thaliana Is Geographically Structured in the Iberian Peninsula. Genetics, 180(2), 1009– 1021. 10.1534/genetics.108.089581

Platt, A., Horton, M., Huang, Y. S., Li, Y., Anastasio, A. E., Mulyati, N. W., Ågren, J., Bossdorf, O., Byers, D., Donohue, K., Dunning, M., Holub, E. B., Hudson, A., Le Corre, V., Loudet, O., Roux, F., Warthmann, N., Weigel, D., Rivero, L., … Borevitz, J. O. (2010). The Scale of Population Structure in Arabidopsis thaliana. PLoS Genetics, 6(2), e1000843. 10.1371/journal.pgen.1000843

Quint, M., Delker, C., Franklin, K. A., Wigge, P. A., Halliday, K. J., & van Zanten, M. (2016). Molecular and genetic control of plant thermomorphogenesis. Nature Plants, 2(1), 15190. 10.1038/nplants.2015.190

R Core Team. (2022). R: A language and environment for statistical computing. R Foundation for Statistical Computing. (4.1.3).

R Core Team. (2025). Support for Parallel Computation in R.

Ristova, D., Carré, C., Pervent, M., Medici, A., Kim, G. J., Scalia, D., Ruffel, S., Birnbaum, K. D., Lacombe, B., Busch, W., Coruzzi, G. M., & Krouk, G. (2016). Combinatorial interaction network of transcriptomic and phenotypic responses to nitrogen and hormones in the Arabidopsis thaliana root. Science Signaling, 9(451). 10.1126/scisignal.aaf2768

Robert J. Hijmans. (2024). raster: Geographic Data Analysis and Modeling.

Robert J. Hijmans. (2025). geodata: Access Geographic Data.

Roeder, A. H. K., Shi, Y., Yang, S., Abbas, M., Sasidharan, R., Yanovsky, M. J., Casal, J. J., Ruffel, S., von Wirén, N., Assmann, S. M., Kinscherf, N. A., Bakshi, A., Alptekin, B., Gilroy, S., SharathKumar, M., Prat, S., & Argueso, C. T. (2025). Translational insights into abiotic interactions: From Arabidopsis to crop plants. The Plant Cell, 37(7). 10.1093/plcell/koaf140

Ruffley, M., Lutz, U., Leventhal, L., Hateley, S., Yuan, W., Keck, J., Rhee, S. Y., Weigel, D., & Exposito-Alonso, M. (2023). Selection constraints of plant adaptation can be relaxed by gene editing. 10.1101/2023.10.16.562583

Satbhai, S. B., Setzer, C., Freynschlag, F., Slovak, R., Kerdaffrec, E., & Busch, W. (2017). Natural allelic variation of FRO2 modulates Arabidopsis root growth under iron deficiency. Nature Communications, 8(1), 15603. 10.1038/ncomms15603

Scheepens, J. F., Deng, Y., & Bossdorf, O. (2018). Phenotypic plasticity in response to temperature fluctuations is genetically variable, and relates to climatic variability of origin, in Arabidopsis thaliana. AoB PLANTS, 10(4). 10.1093/aobpla/ply043

Seren, Ü., Grimm, D., Fitz, J., Weigel, D., Nordborg, M., Borgwardt, K., & Korte, A. (2017). AraPheno: a public database for A. thaliana phenotypes. Nucleic Acids Research, 45(D1), D1054–D1059. 10.1093/nar/gkw986

Subrahmaniam, H. J., Picó, F. X., Bataillon, T., Salomonsen, C. L., Glasius, M., & Ehlers, B. K. (2025). Natural variation in root exudate composition in the genetically structured Arabidopsis thaliana in the Iberian Peninsula. New Phytologist, 245(4), 1437–1449. 10.1111/nph.20314

Sung, S., & Amasino, R. M. (2004). Vernalization in Arabidopsis thaliana is mediated by the PHD finger protein VIN3. Nature, 427(6970), 159–164. 10.1038/nature02195

The 1001 Genomes Consortium, Alonso-Blanco, C., Andrade, J., Becker, C., Bemm, F., Bergelson, J., Borgwardt, K. M., Cao, J., Chae, E., Dezwaan, T. M., Ding, W., Ecker, J. R., Exposito-Alonso, M., Farlow, A., Fitz, J., Gan, X., Grimm, D. G., Hancock, A. M., Henz, S. R., … Zhou, X. (2016). 1,135 Genomes Reveal the Global Pattern of Polymorphism in Arabidopsis thaliana. Cell, 166(2), 481–491. 10.1016/j.cell.2016.05.063

The 1001 Genomes Plus Consortium, Alonso-Blanco, C. C., Ashkenazy, H., Baduel, P., Bao, Z., Becker, C., Caillieux, E., Colot, V., Crosbie, D., De Oliveira, L., Fitz, J., Fritschi, K., Grigoreva, E., Guo, Y., Habring, A., Henderson, I., Hou, X.-H., Hu, Y., Igolkina, A., … Xian, W. (2024). The 1001G+ project: A curated collection of Arabidopsis thaliana long-read genome assemblies to advance plant research. 10.1101/2024.12.23.629943

Thomas Lin Pedersen. (2024a). ggraph: An Implementation of Grammar of Graphics for Graphs and Networks.

Thomas Lin Pedersen. (2024b). tidygraph: A Tidy API for Graph Manipulation.

Togninalli, M., Seren, Ü., Freudenthal, J. A., Monroe, J. G., Meng, D., Nordborg, M., Weigel, D., Borgwardt, K., Korte, A., & Grimm, D. G. (2019). AraPheno and the AraGWAS Catalog 2020: a major database update including RNA-Seq and knockout mutation data for Arabidopsis thaliana. Nucleic Acids Research. 10.1093/nar/gkz925

Toledo, B., Marcer, A., Méndez-Vigo, B., Alonso-Blanco, C., & Picó, F. X. (2020). An ecological history of the relict genetic lineage of Arabidopsis thaliana. Environmental and Experimental Botany, 170, 103800. 10.1016/j.envexpbot.2019.103800

Tonsor, S. J., Scott, C., Boumaza, I., Liss, T. R., Brodsky, J. L., & Vierling, E. (2008). Heat shock protein 101 effects in A. thaliana : genetic variation, fitness and pleiotropy in controlled temperature conditions. Molecular Ecology, 17(6), 1614–1626. 10.1111/j.1365-294X.2008.03690.x

van Eijnatten, A. L., van Zon, L., Manousou, E., Bikineeva, M., Wubs, E. R. J., van der Putten, W. H., Morriën, E., Dutilh, B. E., & Snoek, L. B. (2025). SpeSpeNet: an interactive and user-friendly tool to create and explore microbial correlation networks. ISME Communications, 5(1). 10.1093/ismeco/ycaf036

Vasseur, F., Sartori, K., Baron, E., Fort, F., Kazakou, E., Segrestin, J., Garnier, E., Vile, D., & Violle, C. (2018). Climate as a driver of adaptive variations in ecological strategies in Arabidopsis thaliana. Annals of Botany. 10.1093/aob/mcy165

Wijfjes, R. Y., Boesten, R., Becker, F. F. M., Theeuwen, T. P. J. M., Snoek, B. L., Mastoraki, M., Verheijen, J. J., Güvencli, N., Denkers, L. M., Koornneef, M., van Eeuwijk, F. A., Smit, S., de Ridder, D., & Aarts, M.G. M. (2024). Allelic variants confer Arabidopsis adaptation to small regional environmental differences. The Plant Journal, 120(4), 1662–1681. 10.1111/tpj.17067

Wolfe, M. D., & Tonsor, S. J. (2014). Adaptation to spring heat and drought in northeastern Spanish Arabidopsis thaliana. New Phytologist, 201(1), 323–334. 10.1111/nph.12485

Wu, X., Bellagio, T., Peng, Y., Czech, L., Lin, M., Lang, P., Epstein, R., Abdelaziz, M., Alexander, J., Alonso-Blanco, C., Andersen, H. L., Berbel, M., Bergelson, J., Bossdorf, O., Burghardt, L., Caton-Darby, M., Colautti, R., Delker, C., Dimitrakopoulos, P. G., … Exposito-Alonso, M. (2026). Rapid adaptation and extinction in synchronized outdoor evolution experiments of Arabidopsis. Science, 391(6792). 10.1126/science.adz0777

Yang, W., Cortijo, S., Korsbo, N., Roszak, P., Schiessl, K., Gurzadyan, A., Wightman, R., Jönsson, H., & Meyerowitz, E. (2021). Molecular mechanism of cytokinin-activated cell division in Arabidopsis. Science, 371(6536), 1350–1355. 10.1126/science.abe2305

Yang, W., Feng, H., Zhang, X., Zhang, J., Doonan, J. H., Batchelor, W. D., Xiong, L., & Yan, J. (2020). Crop Phenomics and High-Throughput Phenotyping: Past Decades, Current Challenges, and Future Perspectives. Molecular Plant, 13(2), 187–214. 10.1016/j.molp.2020.01.008

Yim, C., Bellis, E. S., DeLeo, V. L., Gamba, D., Muscarella, R., & Lasky, J. R. (2024). Climate biogeography of Arabidopsis thaliana : Linking distribution models and individual variation. Journal of Biogeography, 51(4), 560–574. 10.1111/jbi.14737

Zan, Y., & Carlborg, Ö. (2019). A Polygenic Genetic Architecture of Flowering Time in the Worldwide Arabidopsis thaliana Population. Molecular Biology and Evolution, 36(1), 141–154. 10.1093/molbev/msy203

Zavafer, A., Bates, H., Mancilla, C., & Ralph, P. J. (2023). Phenomics: conceptualization and importance for plant physiology. Trends in Plant Science, 28(9), 1004–1013. 10.1016/j.tplants.2023.03.023

Ziyatdinov, A., Vázquez-Santiago, M., Brunel, H., Martinez-Perez, A., Aschard, H., & Soria, J. M. (2018). lme4qtl: linear mixed models with flexible covariance structure for genetic studies of related individuals. BMC Bioinformatics, 19(1), 68. 10.1186/s12859-018-2057-x

