## Supplementary materials for "EcoCore; An ecologically diverse panel of *Arabidopsis thaliana* accessions for studying plant-environment interactions"

### Supplemental tables

**Supplemental Table 1: Description of 11 non-redundant environmental parameters used to define environmental groups of *A. thaliana* accessions.** Indicated are the variable name as listed in AraClim, variable interpretation (description), and data source.

| Cluster # | Variable | Description | Source |
|---|---|---|---|
| 1 | CRU_Tmp_summer | Daily mean temperature summer | CRU (Climatic Research Unit) |
| 2 | MERRAclimmean_90s_bio15 | Mean precipitation seasonality in 1990s | Modern Era Retrospective Analysis for Research and Applications Reanalysis (MERRA clim) |
| 3 | MERRAclimmean_00s_bio16 | Mean precipitation wettest quarter in 2000s | Modern Era Retrospective Analysis for Research and Applications Reanalysis (MERRA clim) |
| 4 | MERRA_climmin_00s_bio9 | Minimum mean temperature driest quarter in 2000s | Modern Era Retrospective Analysis for Research and Applications Reanalysis (MERRA clim) |
| 5 | MERRA_climmax_90s_bio3 | Maximum isothermality in 1990s | Modern Era Retrospective Analysis for Research and Applications Reanalysis (MERRA clim) |
| 6 | MERRA_climmax_00s_bio2 | Maximum mean diurnal range in 2000s | Modern Era Retrospective Analysis for Research and Applications Reanalysis (MERRA clim) |
| 7 | CHELSA_BIO13 | Mean precipitation wettest month 1979 – 2013 | CHELSA (Climatologies at high resolution for the earth's land surface areas) |

| | | | |
|---|---|---|---|
| 8 | CHELSA_BIO7 | Mean temperature annual range 1979 – 2013 | CHELSA (Climatologies at high resolution for the earth's land surface areas) |
| 9 | FAO_Aridity_index_de_Martonne_GPCC_Fulldata | Mean aridity 1976 – 2000 | FAO (Food and Agriculture Organization) |
| 10 | ODIAC_Fossil_fuel_emissions_spring | Mean fossil fuel emissions in spring 2000 – 2009 | ODIAC (Open-source Data Inventory for Anthropogenic CO2) |
| 11 | CO_Summer (MOPITT) | Mean carbon monoxide emissions summer 2001 – 2010 | MOPITT (Measurements of Pollution in the Troposphere) |

### Supplemental figures

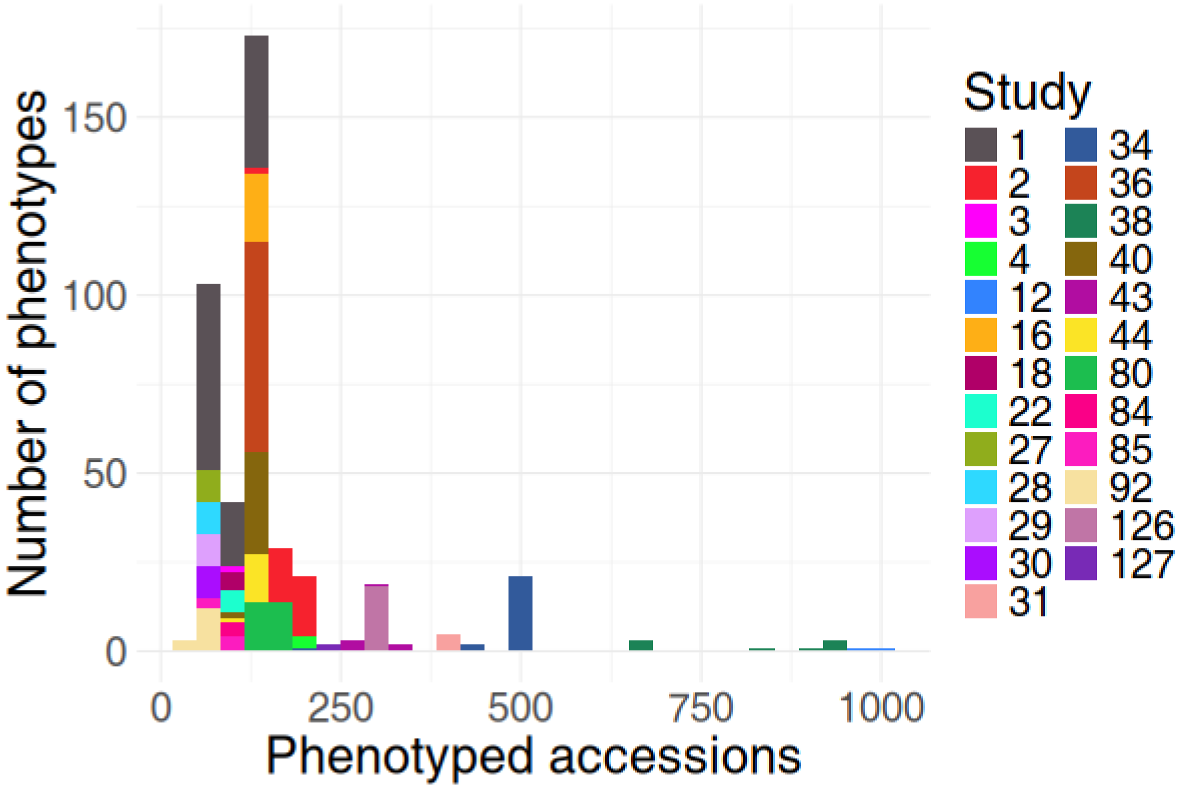


**Supplemental Figure 1: Density distribution of real panel sizes used in studies included in the AraPheno database.** Indicated is the number of accessions from the 1001G panel included per trait. Colors represent the individual studies (reference number corresponding to the AraPheno database entry, see also **Supplemental table 1**).

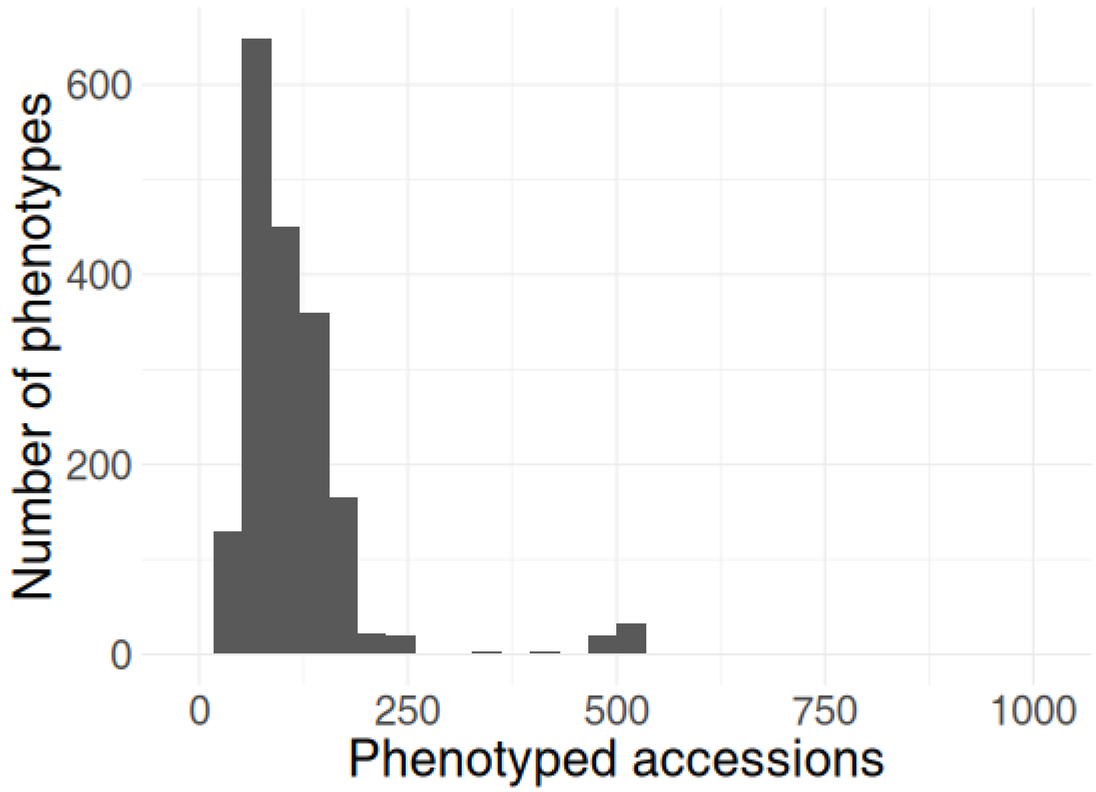


**Supplemental Figure 2: Number of accessions phenotyped per trait in an extension of the AraPheno database.** The extended dataset contained a total of 1863 traits (Ruffley et al., 2024). The median number of phenotyped accessions was 102 accessions compared to 130 in the AraPheno database.

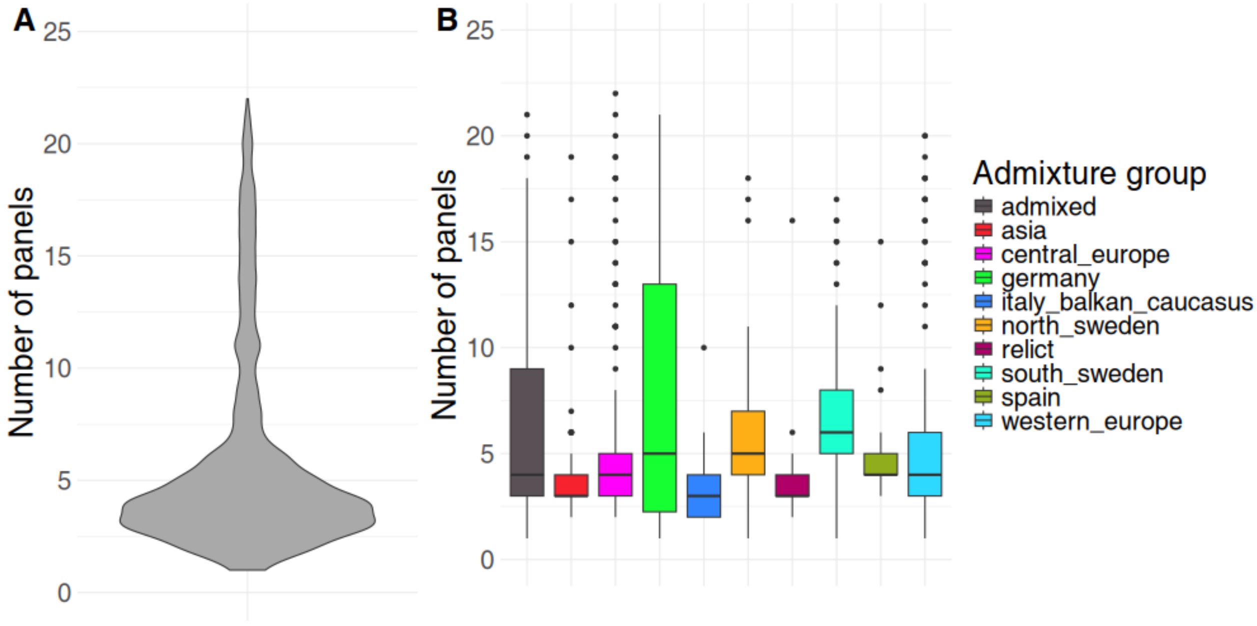


**Supplemental Figure 3: Accession frequency in the AraPheno database.** Distribution of **A**) the number of times each individual 1001G accession and **B)** different admixture groups (derived from The 1001 Genomes Consortium, 2016), are included in a sub-panel present in the AraPheno database for at least one trait.

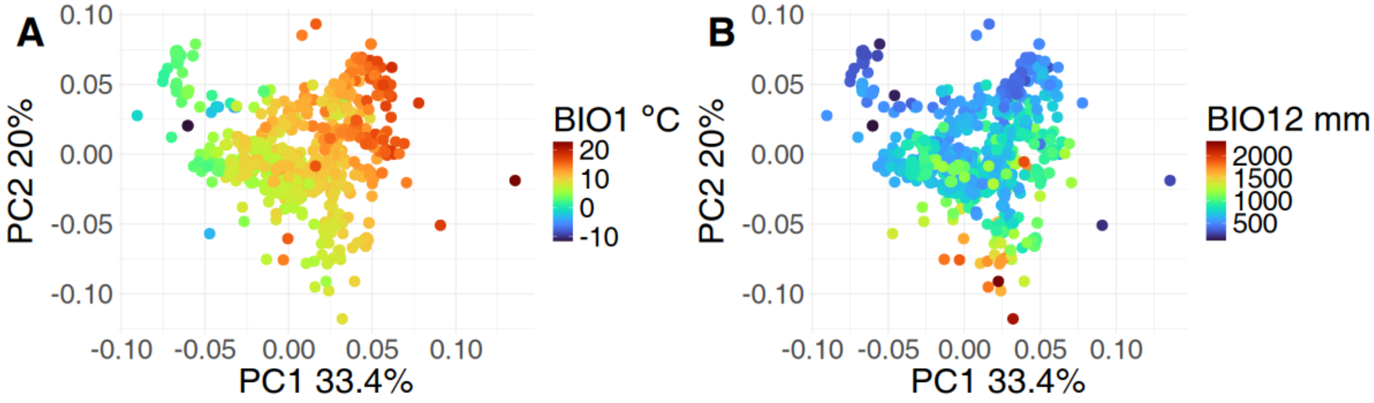


**Supplemental Figure 4: PCA analysis on variation in AraClim v2 entries across accessions.** A total of 1095 accessions and 373 numeric environmental variables are included. Accessions are projected onto the first two principal components. Accessions colored by **A)** BIO1 (mean annual temperature (°C)) and **B**) BIO12 (Annual precipitation (mm)).

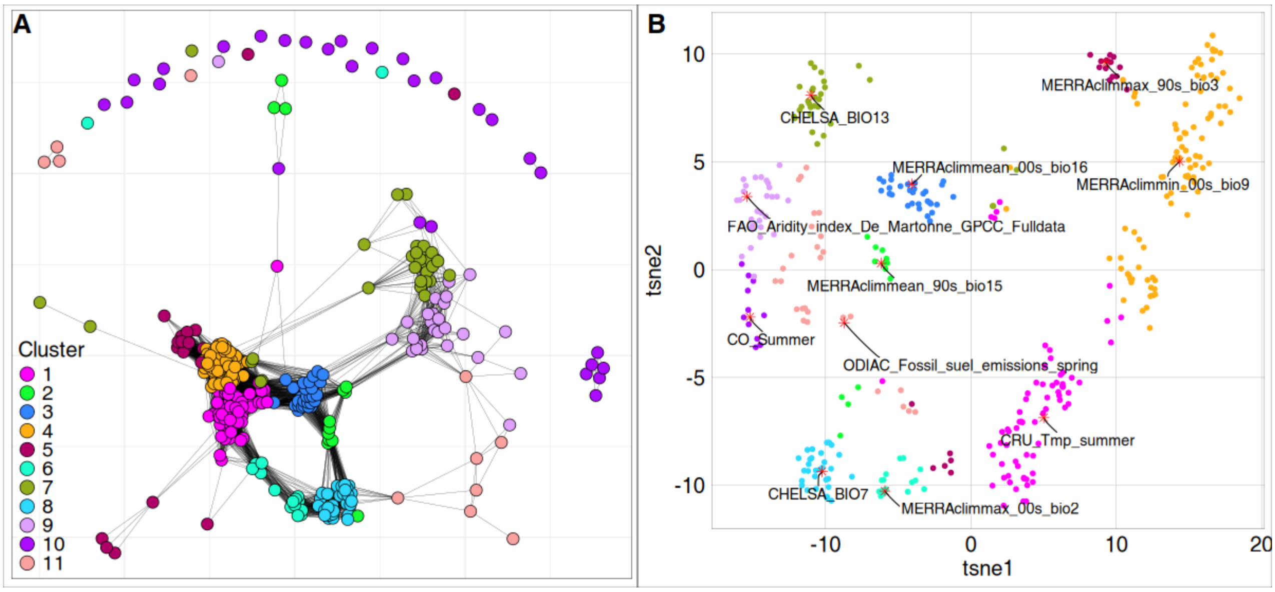


**Supplemental Figure 5: Redundancy among environmental parameter values of the AraClim database. A)** Correlation network wherein each node is an environmental parameter and each edge is a Pearson correlation > 0.6. Correlations are based on all accessions in the 1001G panel. Environmental parameters with more than 10 NA's were removed prior to clustering and correlation network construction. Colors indicate the cluster assignment of a k-means clustering (K = 11) done on the 373*373 correlation matrix of the environmental parameters in AraClim. **B)** t-SNE analysis on mean-centred and SD-normalized AraClim database environmental parameters. Each dot represents an environmental variable and the color shows the cluster assignment of the same k-means clustering as in panel A. Red stars and text show the environmental parameters closest to the centroid of each of the 11 k-means clusters. These environmental parameters were used to design environmental groups of accessions.

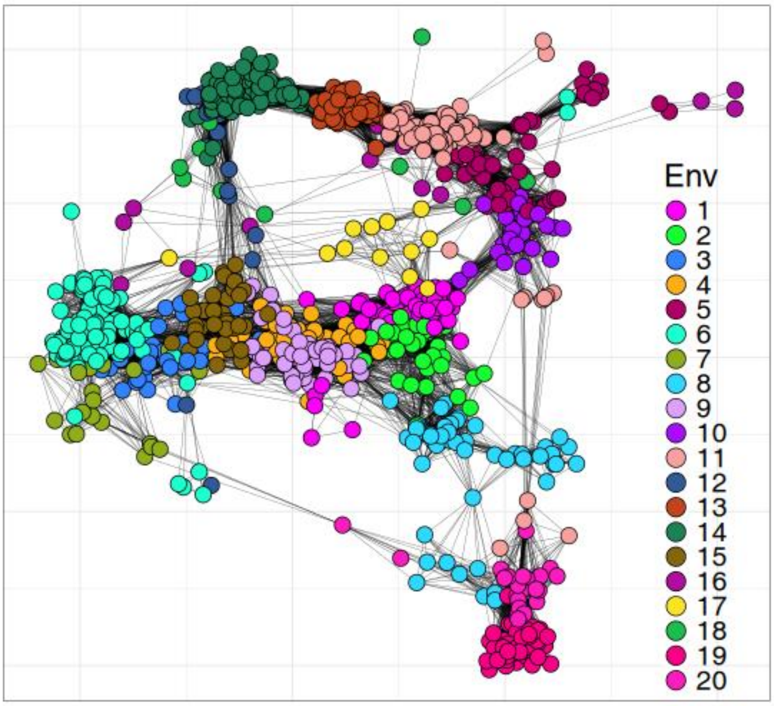


**Supplemental Figure 6: Environmental correlations between accessions.** Correlation network where each node is an accession and each edge is a Pearson correlation > 0.75 based on 11 non-redundant environmental parameters. The color indicates the environmental group assignment.

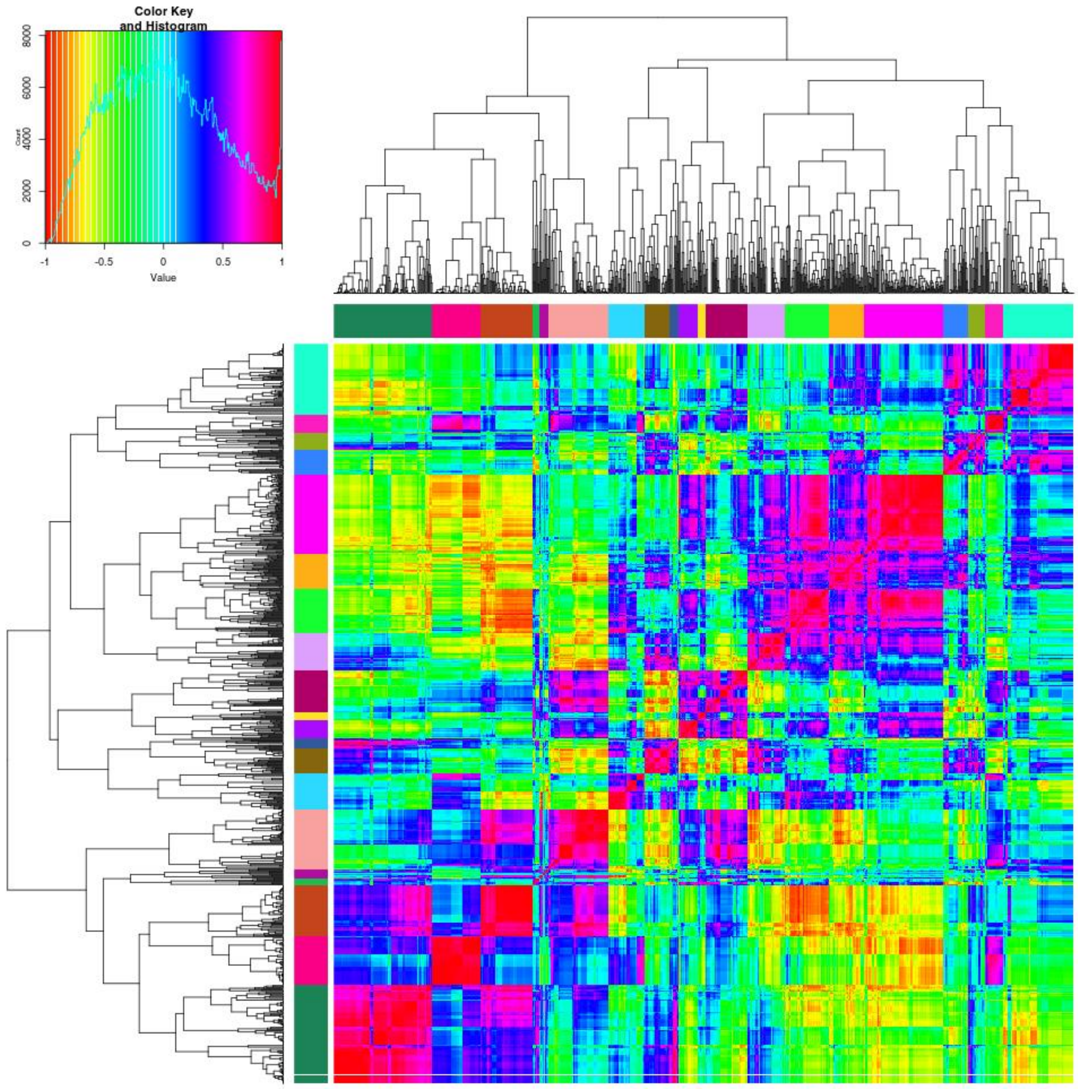


**Supplemental Figure 7: Hierarchical clustering of correlations between accessions based on the 11 non-redundant variables shown in table 1.** 1066 accessions from the 1001G were clustered into 20 environmental groups (row and column colors) by cutting the dendrogram to 20 groups**.** Colors in heatmap indicate Pearson correlation values.

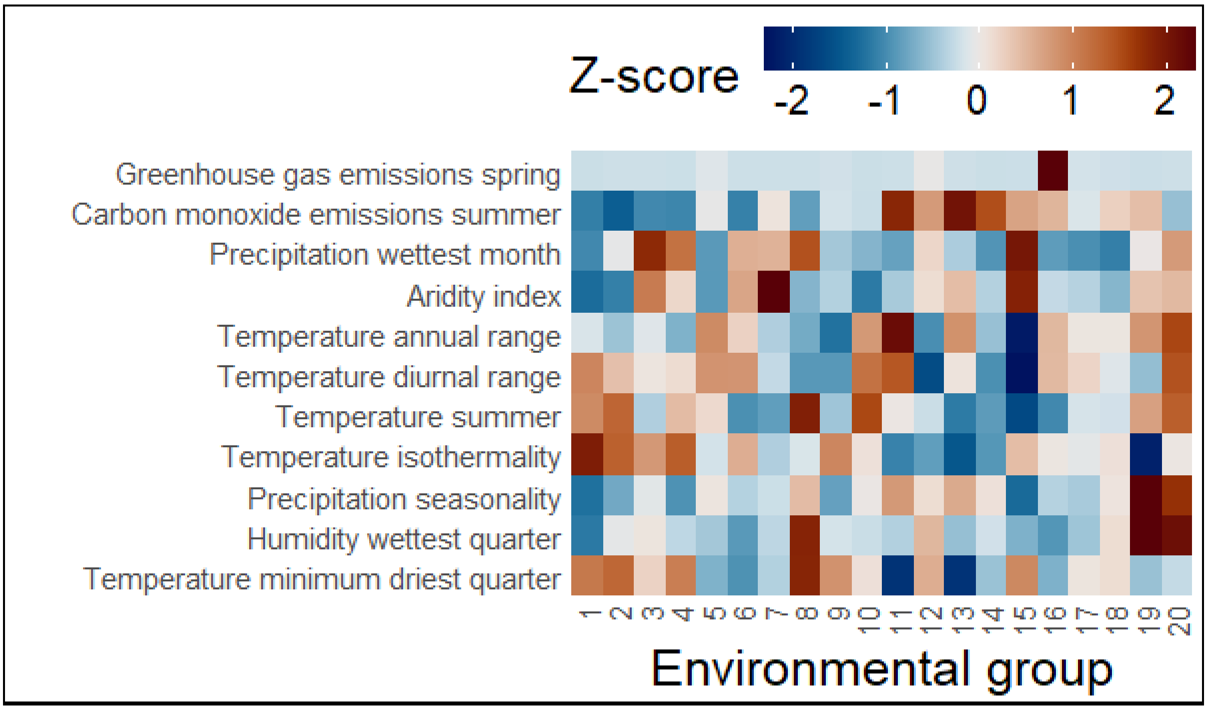


**Supplemental Figure 8: Mean z-score per environmental group for the 11 non-redundant variables used for constructing the 20 environmental groups.** Heatmap showing for each environmental group the relative value expressed as z-score for worlclim variables obtained from the AraClim v2 database.

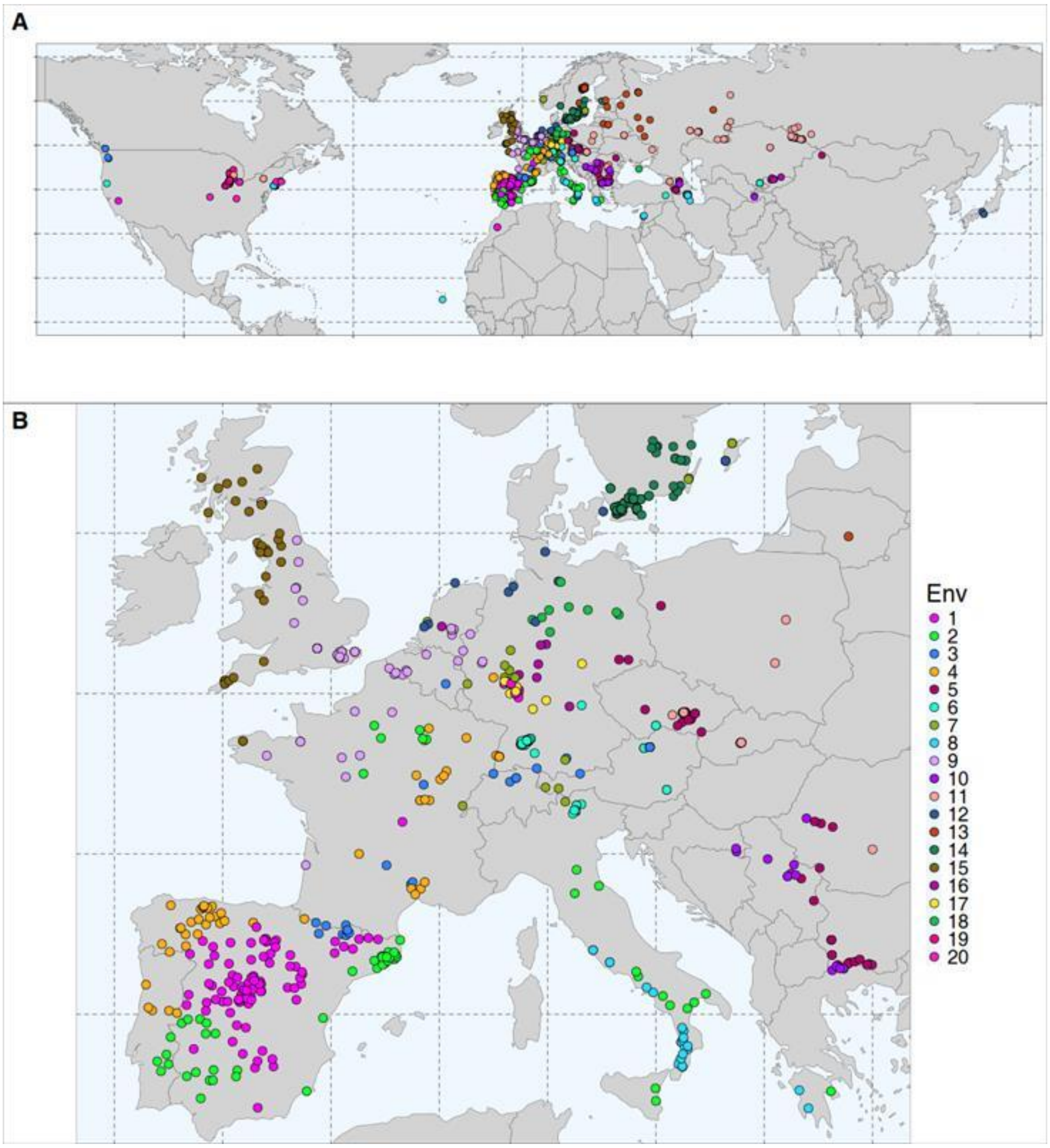


**Supplemental Figure 9: Geographic distribution of 1001G accessions per environmental group. A)** World map, showing the geographic origins of the 1001G accessions color-coded per environmental group and **B)** zoom-in on Europe.

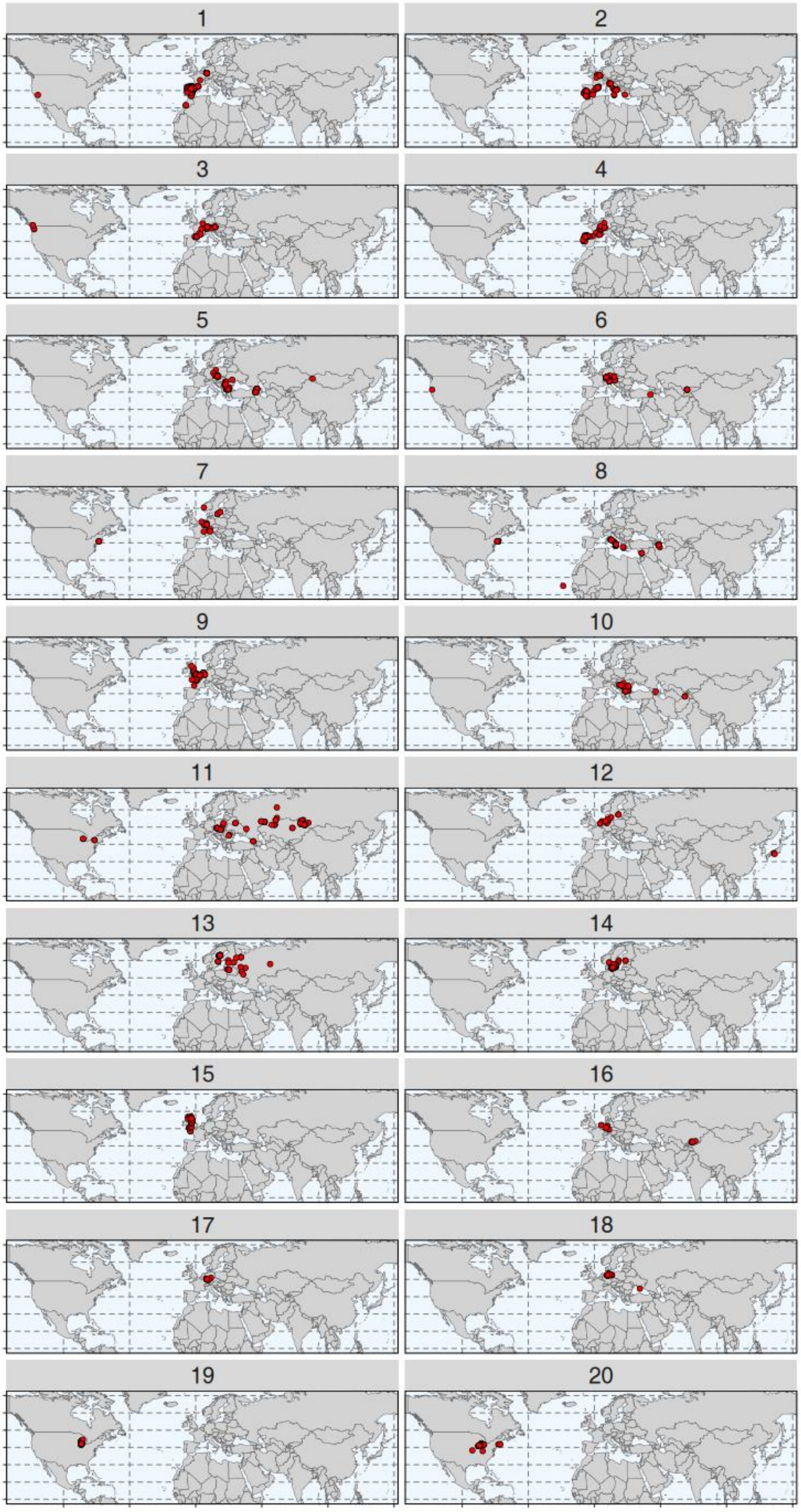


**Supplemental Figure 10: Geographic distribution of accessions per environmental group. A)** world maps with red dots showing the geographic origin of accessions per environmental group (environmental group number indicated above each panel).

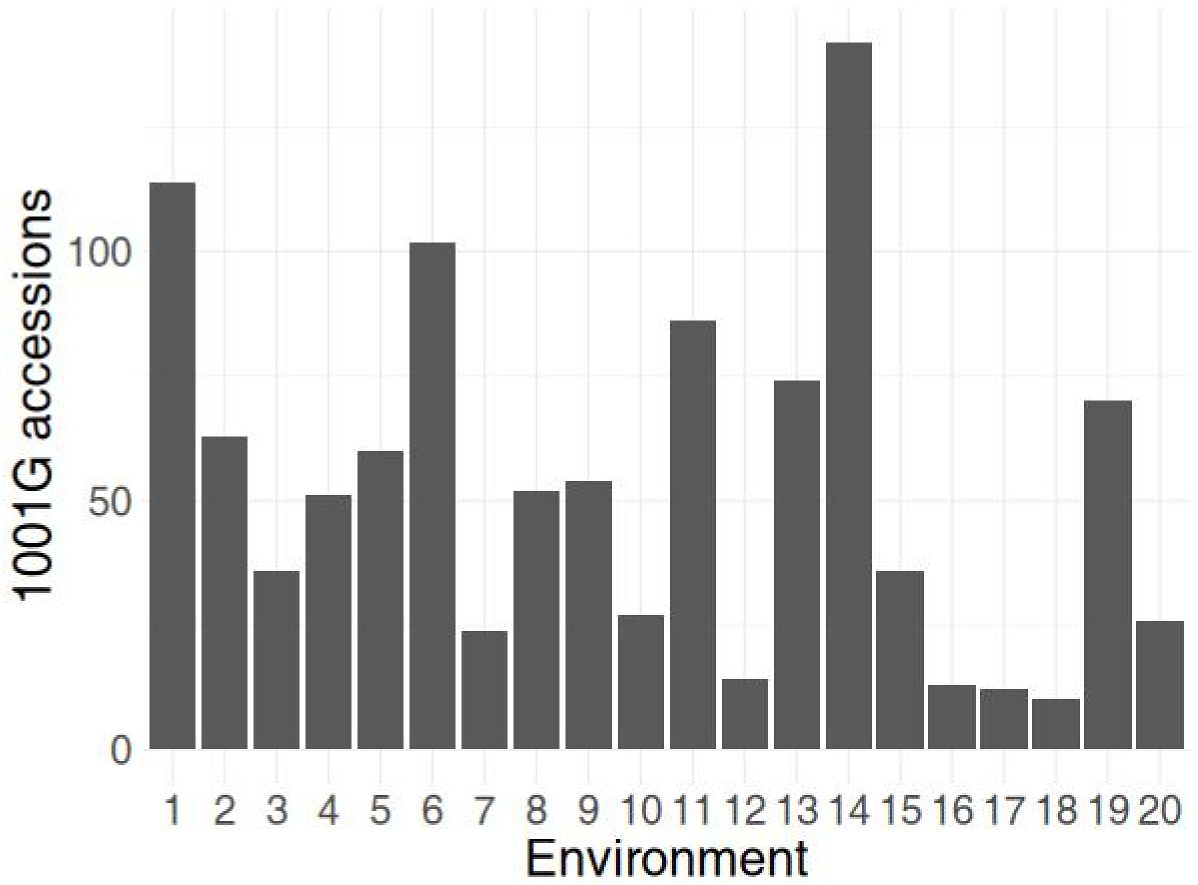


**Supplemental figure 11: Number of 1001G individual accessions per assigned environmental group.**

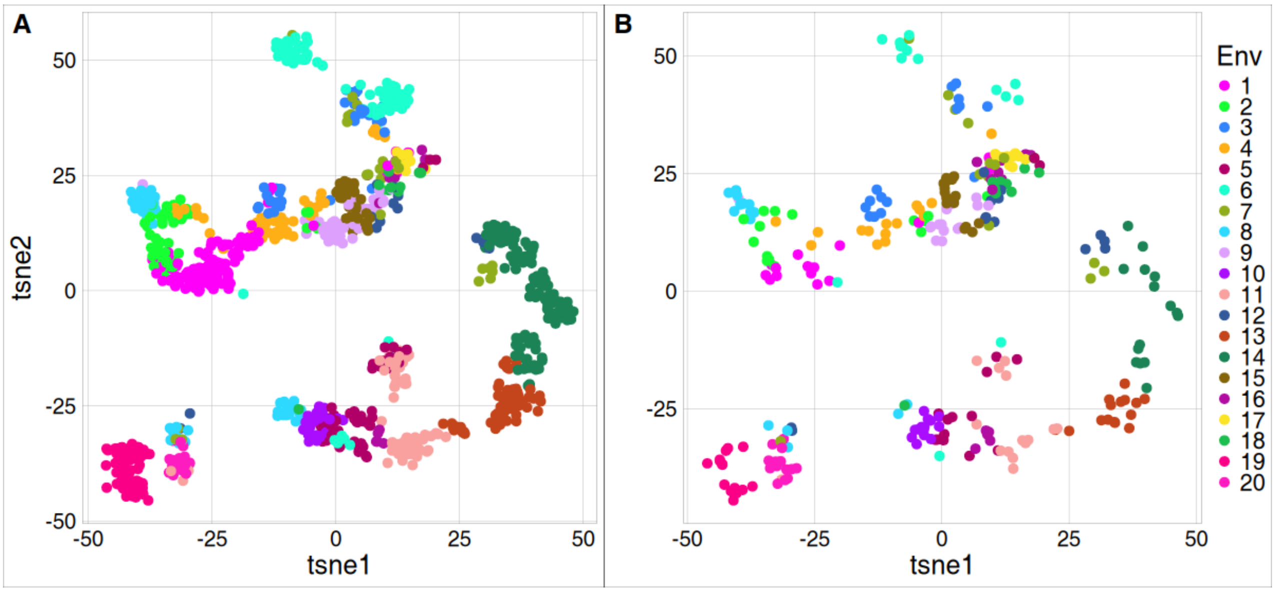


**Supplemental Figure 12: t-SNE analysis on the full AraClim database (373 variables).** Each dot represents an accession of **A**) the 1001G panel accessions and **B**) the EcoCore panel. Colors indicate the environmental group.

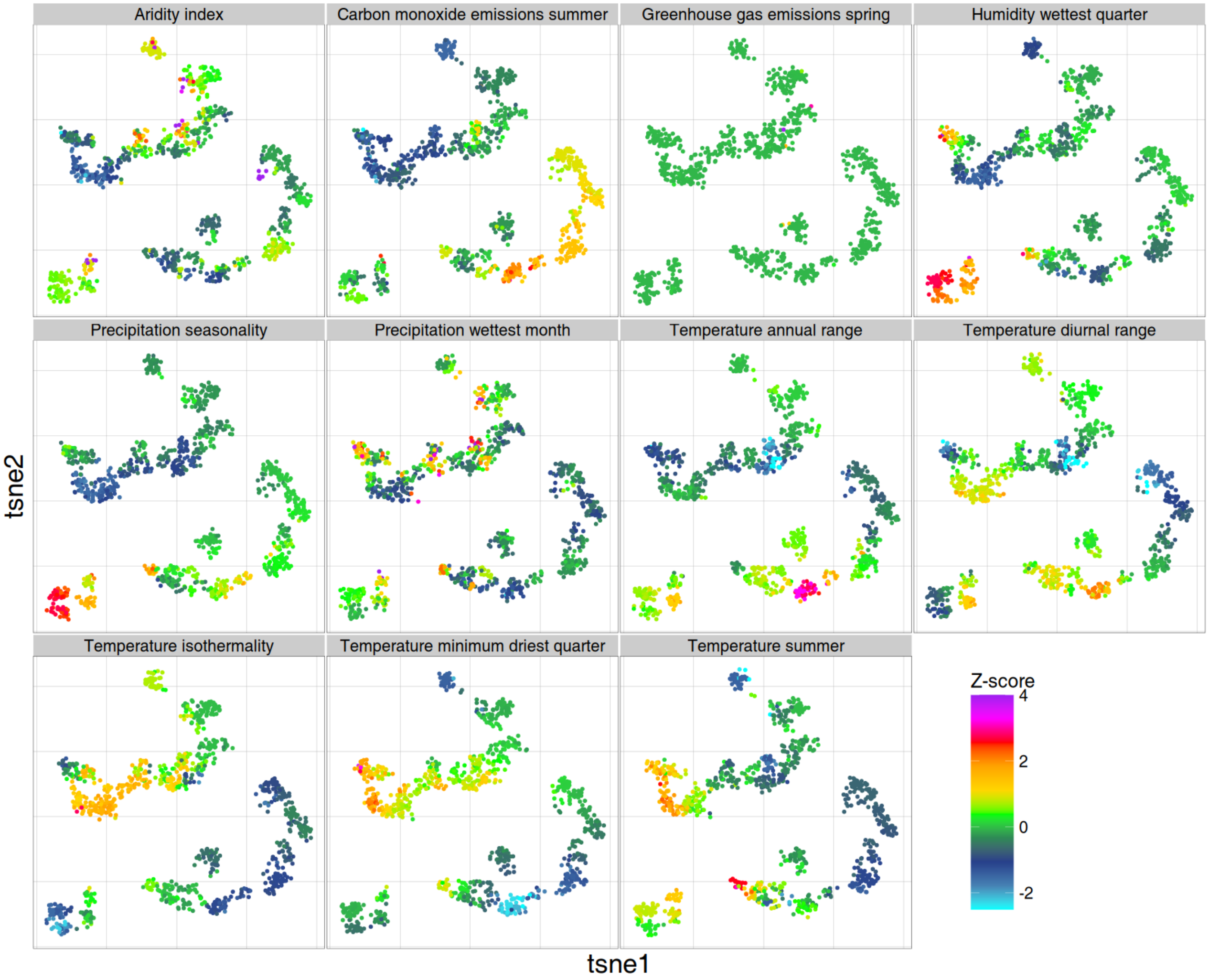


**Supplemental Figure 13: Relationship between t-SNE visualization on the full AraClim database (373 variables) and the 11 K-mean analysis-derived non-redundant variables.** Same t-SNE as in Supplemental Figure 12 with each dot representing an accession. Colors indicates the z-score for each of the cluster-representative environmental parameters (indicated above each panel) derived using k-means.

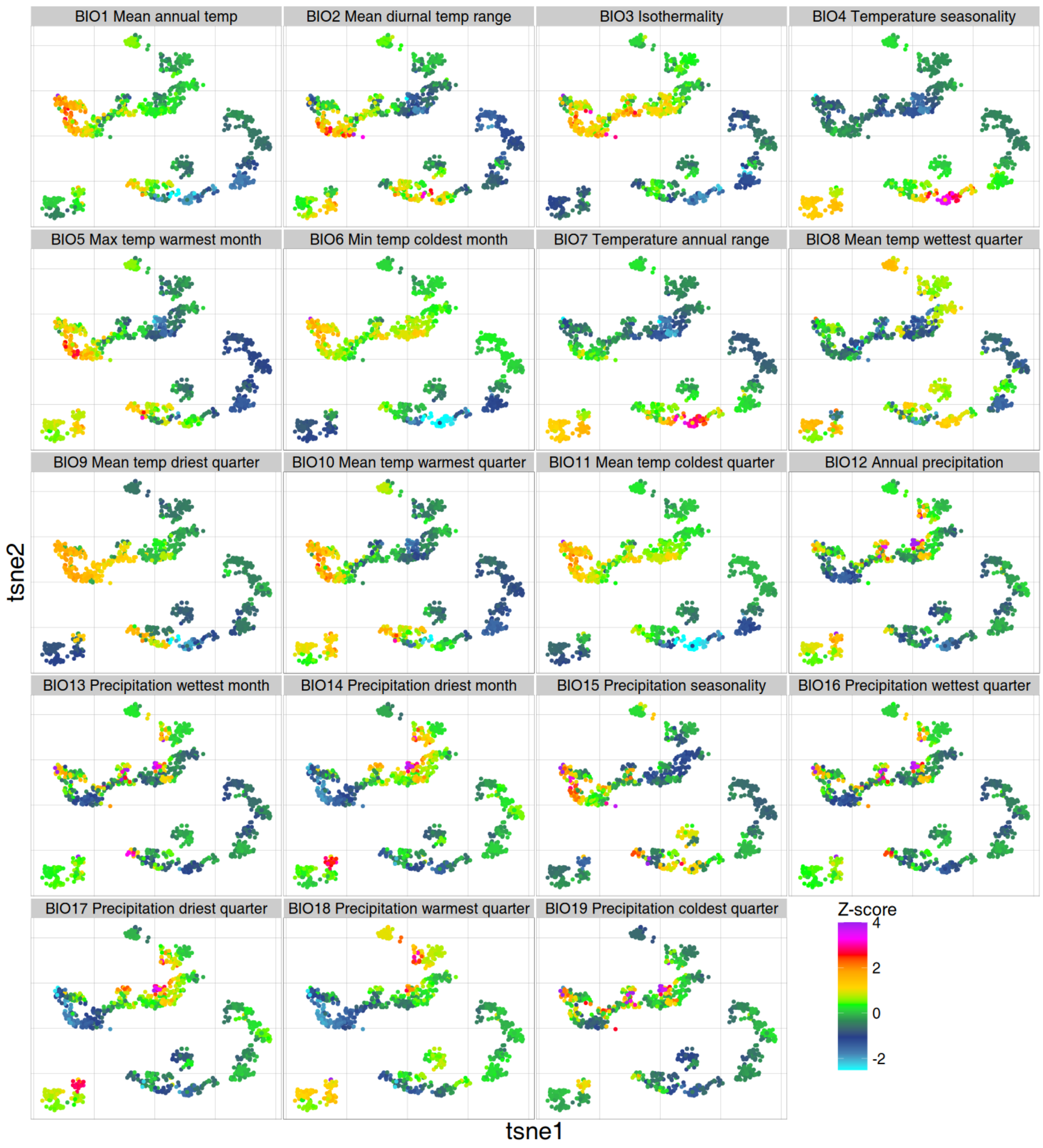


**Supplemental figure 14: t-SNE analysis on the full AraClim database (373 variables), showing the relationship with the 19 WorldClim 2 variables.** Same t-SNE as in Supplemental Figure 12 and Supplemental Figure 13 with each dot representing an accession. Colors indicate the z-score for each WordClim 2 parameter (indicated above each panel).

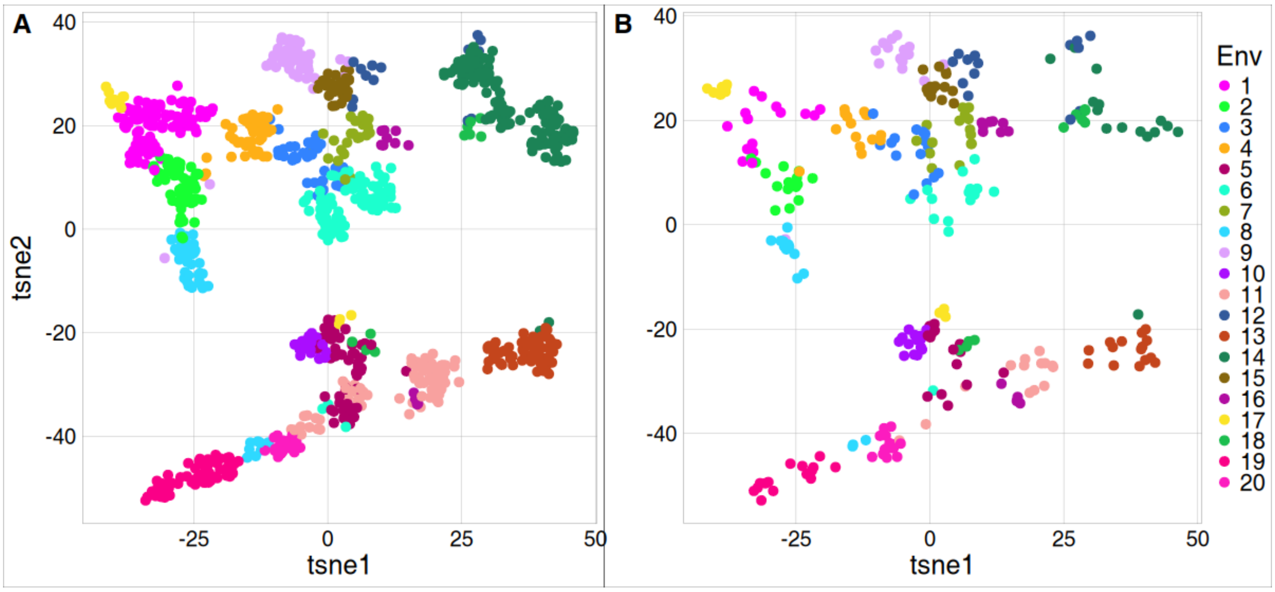


**Supplemental Figure 15: t-SNE analysis on the 11 K-mean analysis-derived non-redundant environmental variables.** Each dot represents an accession of **A)** all 1001G accessions and **B)** the EcoCore panel. Colors indicate the environmental group.

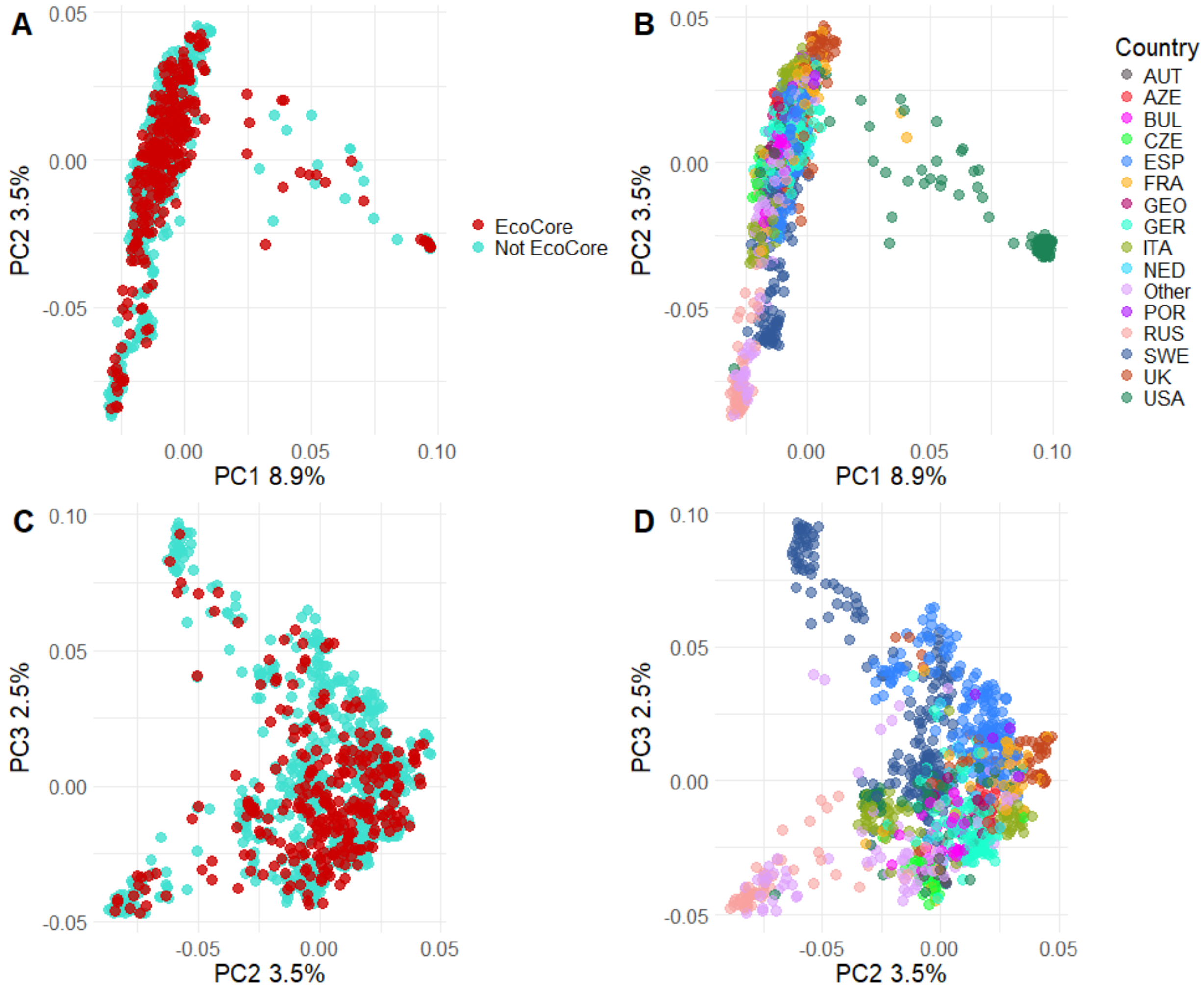


**Supplemental Figure 16: PCA of a high-density SNP variant map of the full 1001G panel.** For the PCA all variants with a MAF > 30% (412489 variants for 1066 accessions) were used (Arouisse et al., 2020). **A)** Genetic variation over PC1 and PC2. Blue dots represent 1001G accessions and red dots represent accessions selected for the EcoCore panel. **B)** Same plot as in A), showing the country of origin of the accessions. **C)** Genetic variation over PC2 and PC3. Blue dots represent 1001G accessions and red dots represent accessions selected for the EcoCore panel**. D)** Same plot as in C), showing the country of origin of the accessions.

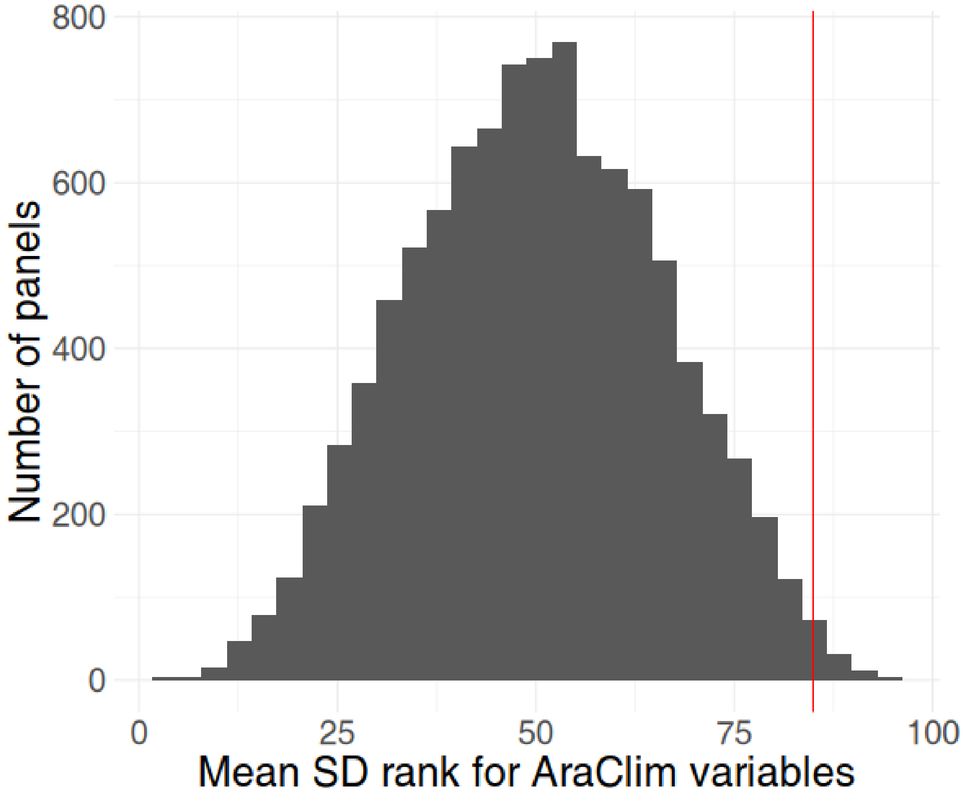


**Supplemental Figure 17: Mean SD rank of the EcoCore panel compared to random panels for the AraClim variables.** Red line shows the mean rank of the EcoCore panel, compared to the distribution of the mean rank over all AraClim variables (grey) for 10000 generated random panels of the same size as the EcoCore panel.

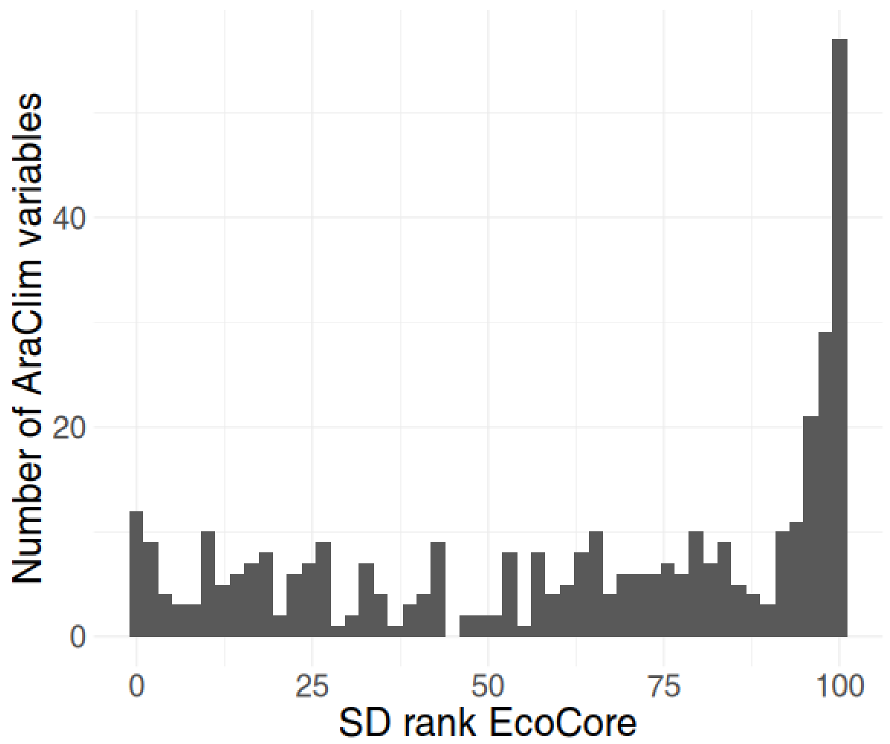


**Supplemental Figure 18: Distribution of SD percentile of the EcoCore panel across AraClim environmental variables.** Indicated is the percentile of the EcoCore panel relative to 10000 random generated panels of the same size over all 373 AraClim variables. A higher value on the x-axis means that the SD is high compared to the random panels.

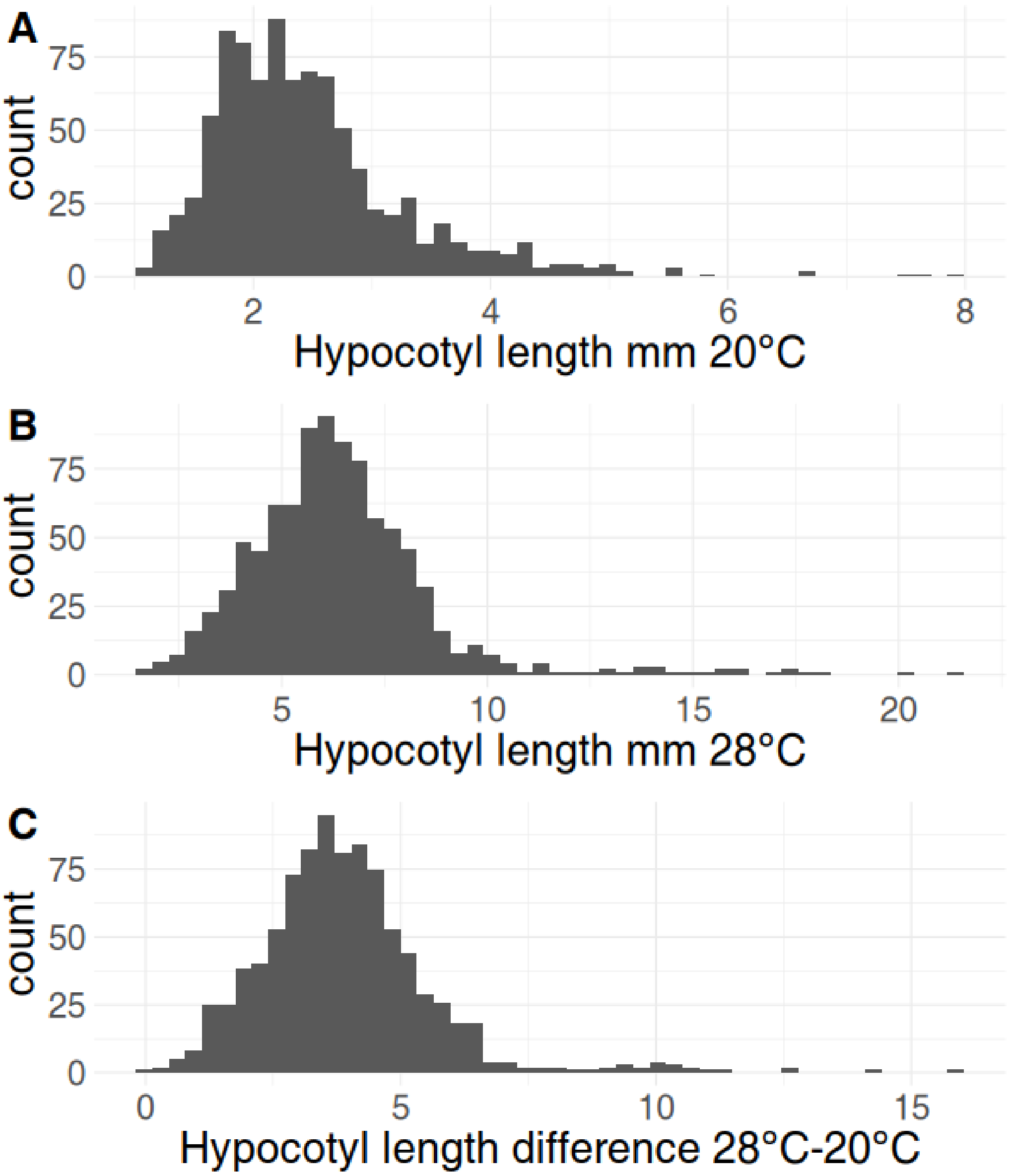


**Supplemental Figure 19: Distribution of hypocotyl lengths of 913 accessions of the 1001G panel. A)** In 20$^{o}$C. **B)** In 28$^{o}$C. **C)** Difference in hypocotyl lengths between the two temperature conditions (28$^{o}$C-20$^{o}$C).

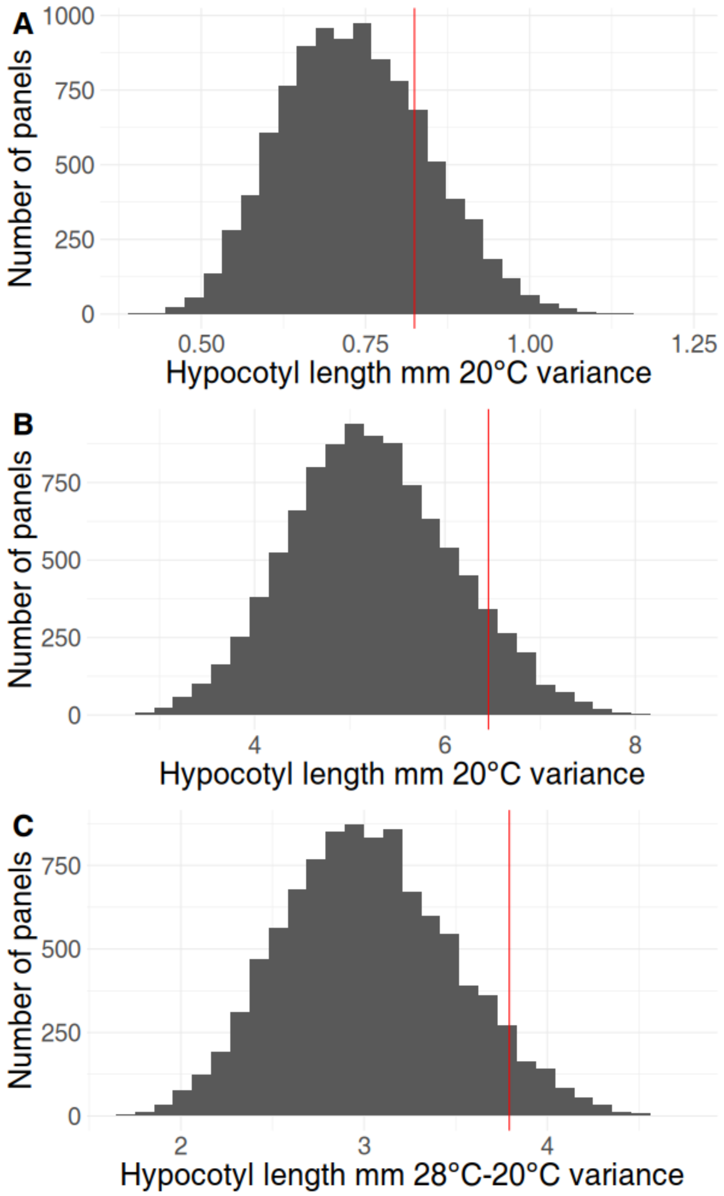


**Supplemental Figure 20: Variance in hypocotyl lengths in random panels compared to the EcoCore panel.** Histograms indicate distribution of the hypocotyl length variance in 9999 generated random panels. Red vertical lines indicate variance of the EcoCore panel for hypocotyl length in **A)** 20$^{o}$C, **B)** 28$^{o}$C, and **C)** the difference in hypocotyl lengths (28$^{o}$C-20$^{o}$C).

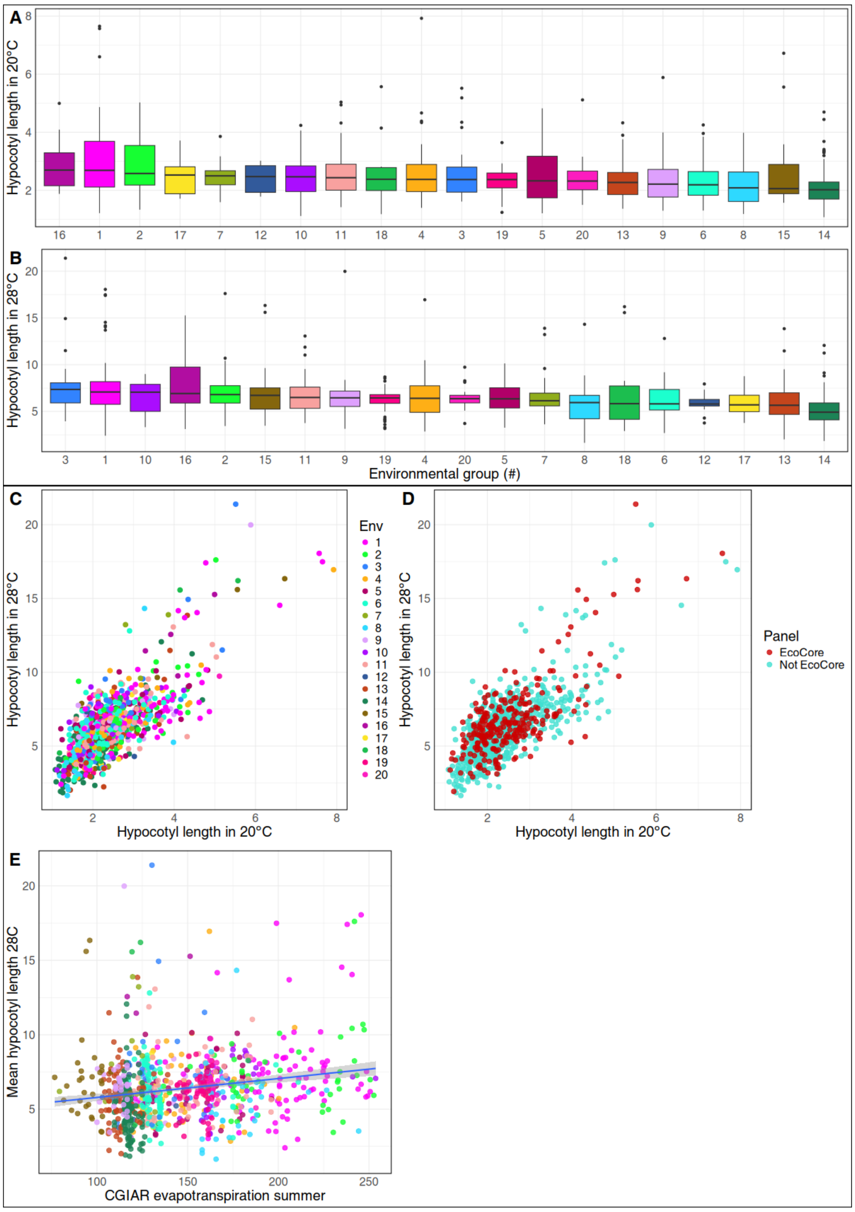


**Supplemental Figure 21: Relationship between hypocotyl lengths and local environment of collection.** Distribution per environmental group of the hypocotyl length **A)** in 20ºC and **B** in 28ºC, for 913 available accessions of the 1001G panel. Environmental groups are ordered by the median hypocotyl length. **C, D)** Relationship between hypocotyl length in 20ºC and 28 ºC. **C)** Colors correspond to the 20 assigned environmental groups. **D)** Blue dots are 1001G accessions, and red dots are accessions selected for the EcoCorepanel. **E)** Relationship between the evapotranspiration predicted by Consultative Group on International Agricultural Research (CGIAR) (obtained from AraClim) and the hypocotyl length in 20 ºC. Colors correspond to the 20 assigned environmental groups.

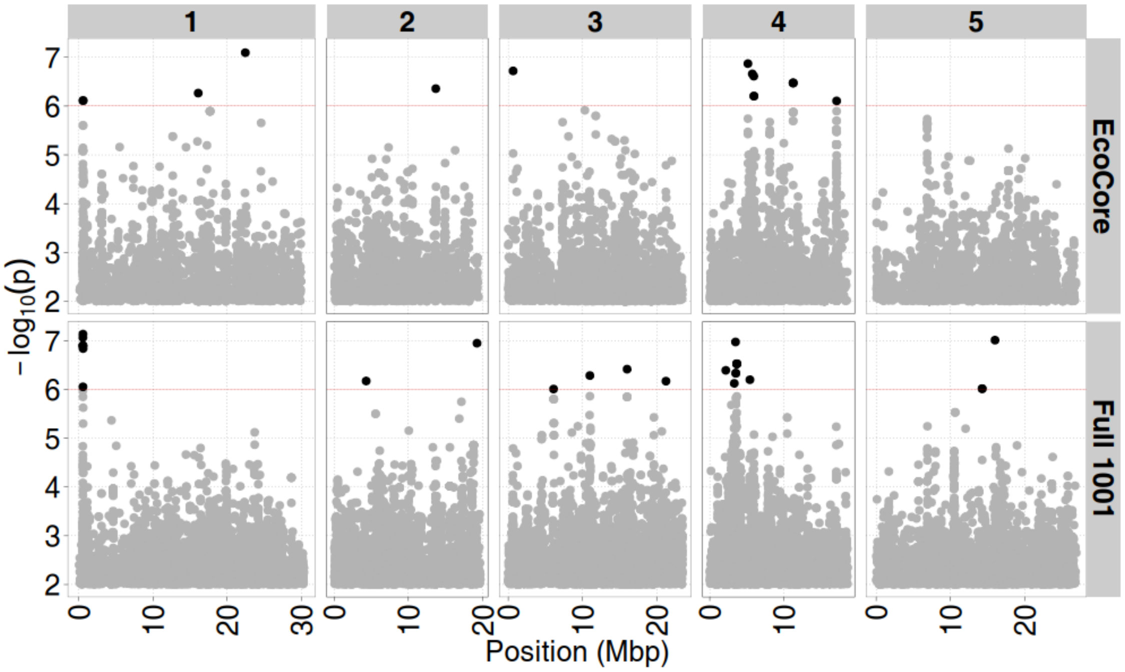


**Supplemental Figure 22: GWAS results for hypocotyl length in 28°C.** Indicated are associations mapped with **A)** the 1001G panel (966 accessions) and **B)** mapped with the EcoCore panel (256 accessions). $-\log_{10}(p)$ indicates the significance of the mapped genetic variants across the 5 chromosomes of Arabidopsis (genetic position indicated in mega bases pairs; Mbp). The red line indicates the threshold of significance, above which significant associations are indicated in bold.

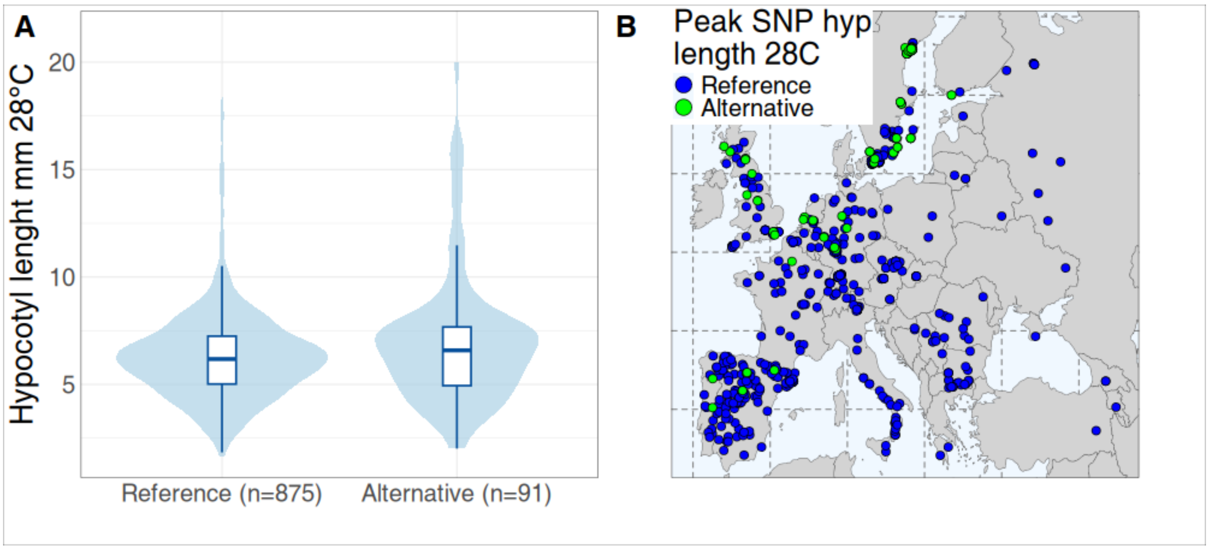


**Supplemental Figure 23: Effect size and geographic distribution of allele associated with hypocotyl length at 28ºC**. Variant genome position is chromosome 1; 0.585122 Mbp. **A)** Distribution of hypocotyl lengths in 28ºC separated by allele variant. **B)** Map showing all European 1001G accessions. Color indicates allele (blue = reference, green = alternative allele). No accessions with the alternative allele are present outside of Europe within the 1001G.

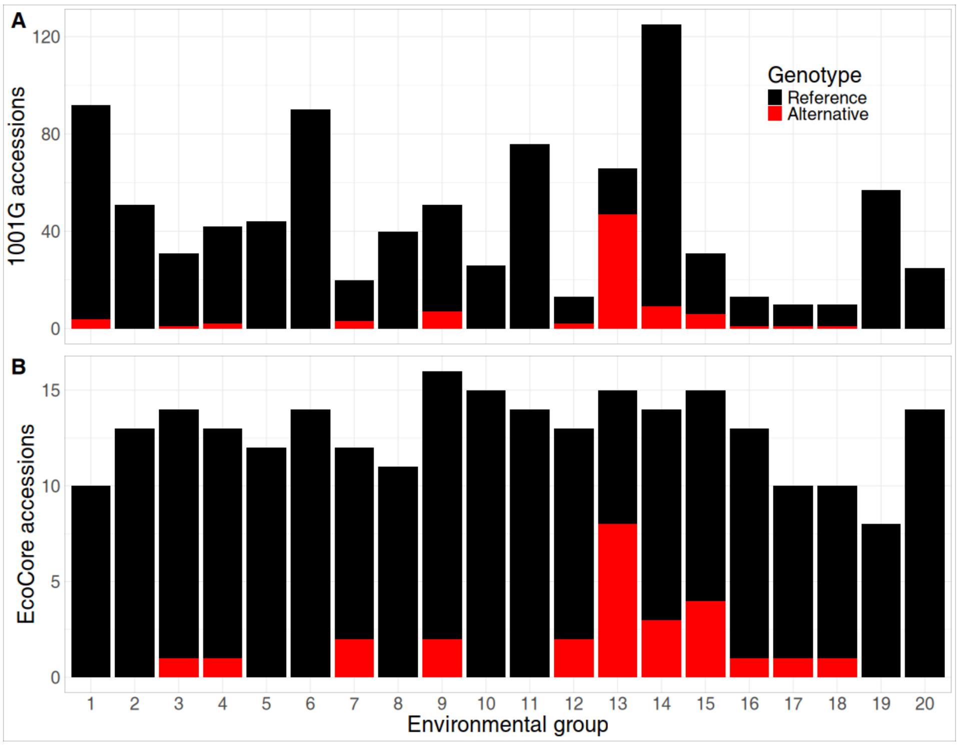


**Supplemental Figure 24: Allele distribution per environmental group of a genetic variant associated with hypocotyl length at 28°C**. Variant genome position is chromosome 1; 0.585122 Mbp. **A)** 1001G panel. **B)** EcoCore panel. Black = reference allele, red = alternative allele.

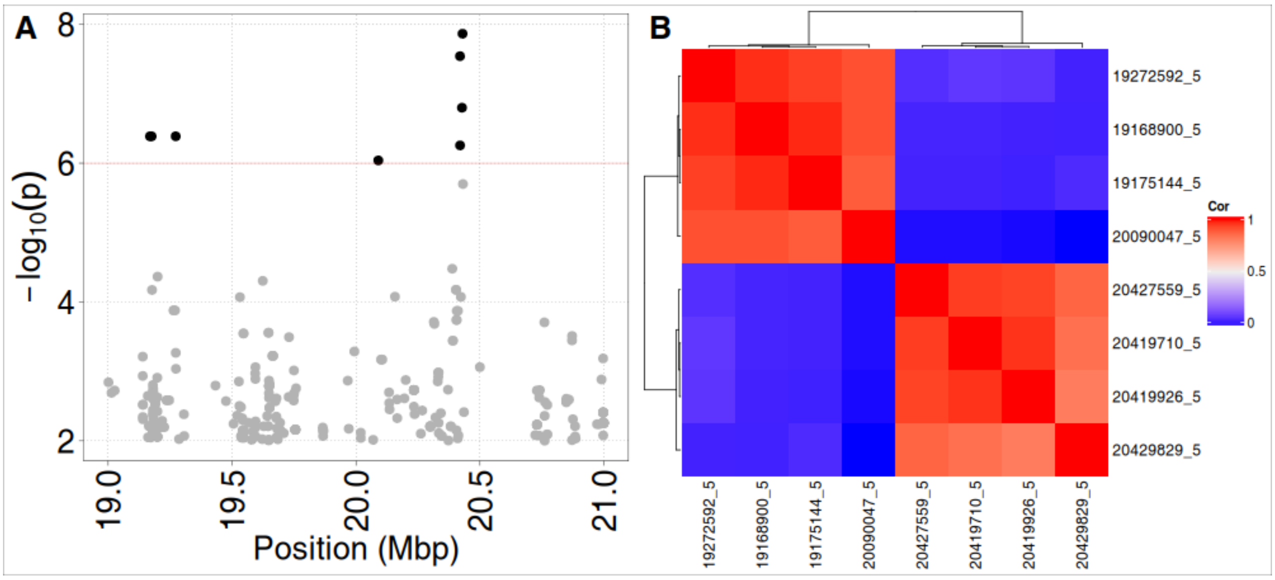


**Supplemental figure 25: Zoom-in on QTL for hypocotyl length in 20°C on chromosome 5 between 19 Mbp and 20.5 Mbp. A)** GWAS results. $\log_{10}$(p) indicates the significance of the mapped genetic variants, genetic position indicated in mega bases pairs; Mbp) **B)** Pearson correlations between the significant SNPs visible in panel A) based on their alleles.

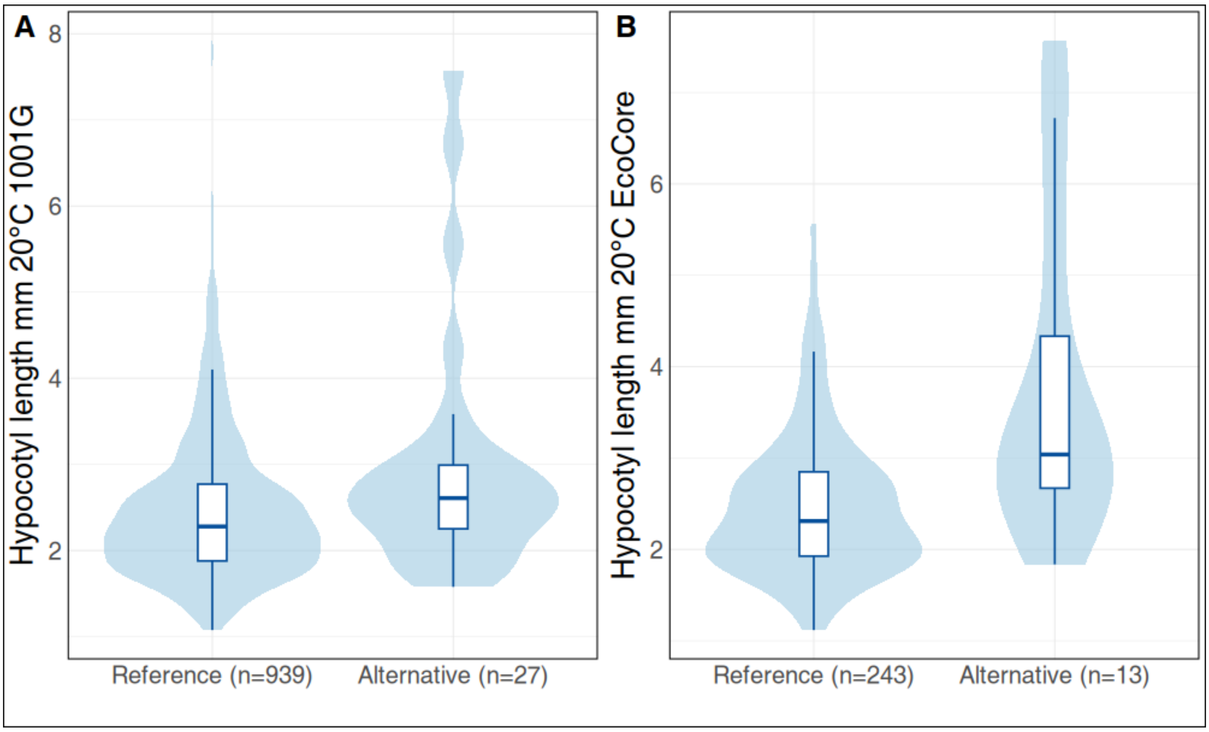


**Supplemental Figure 26: Effects size of chromosome 5 peak allele associated with hypocotyl length at 20ºC**. Variant genome position is chromosome 5; 20.429829 Mbp, see **Figure 4**. Distribution in **A)** The full 1001G panel and **B)** the EcoCore panel.

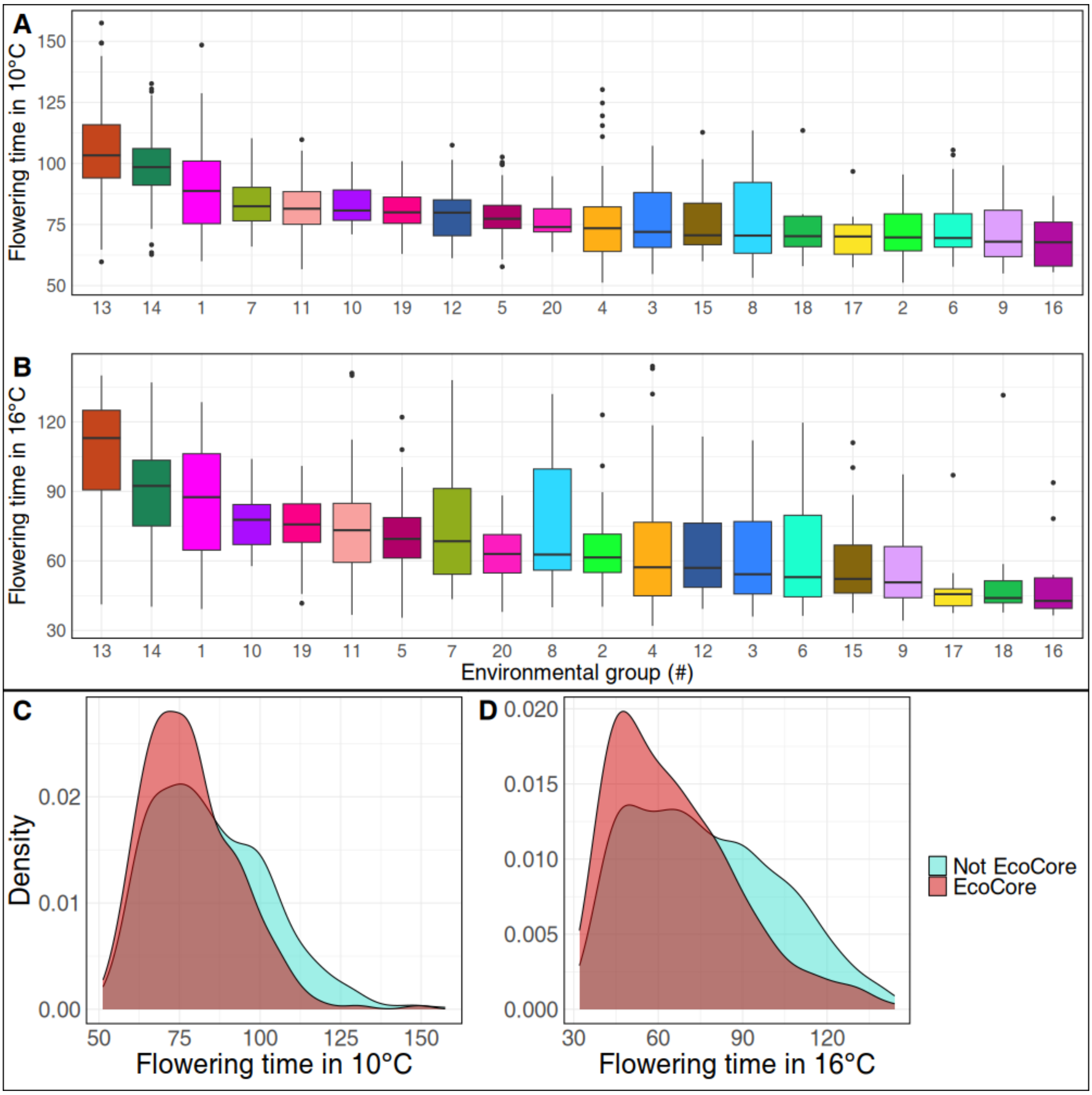


**Supplemental Figure 27: Relationship between flowering time and local environment in the 1001G and EcoCore panels.** Distribution of flowering time in **A)** 10°C and **B)** 16°C per environmental group, for A) 943 and B) 916 available accessions of the 1001G panel. Environmental groups are ordered by the median flowering time. Distribution of flowering time in **C)** 10 °C and **D)** 16 °C in the 1001G, split by whether accessions are part of the EcoCore (red) or not (blue).
